# Impact of inter-species hybridisation on antifungal drug response in the *Saccharomyces* genus

**DOI:** 10.1101/2024.01.29.577396

**Authors:** Federico Visinoni, William Royle, Rachel Scholey, Yue Hu, Soukaina Timouma, Leo Zeef, Edward J Louis, Daniela Delneri

## Abstract

Antifungal drug resistance across fungal and yeast pathogens presents one of the major concerns for global public health. Understanding the interactions between genetic background and environment is important for the development of new, effective treatments of infections. Allelic variation within populations of Ascomycota as well as hybridisation impacts the phenotype in response to stressful conditions, including to antifungal drugs. We exploited recent advances in multigenerational breeding of *Saccharomyces* interspecies hybrids to study the impact of hybridisation on antifungal resistance and identify quantitative trait loci (QTL) responsible for the phenotypes observed. A library of *Saccharomyces cerevisiae* x *S. kudriavzevii* hybrid offspring was screened in the presence of sub-lethal concentrations of six antifungal drugs and revealed a broad phenotypic diversity across the progeny. QTL analysis was carried out comparing alleles between the pools of high and low fitness offspring, identifying hybrid-specific genetic regions involved in resistance to fluconazole, micafungin and flucytosine. We found both drug specific and pleiotropic regions, and through gene ontology and SIFT analysis we identify potential causal genes, such as *BCK2* and *DNF1* that were validated via reciprocal hemizygosity analysis. We highlight 41 regions that contain genes not previously associated with resistance phenotypes in the literature. The results of this screening will help identify new pathways contributing to drug resistance, and lead to greater understanding of how allelic variation, hybridisation and evolution affect antifungal drug resistance in yeast and fungi.

## Introduction

The tally of life-threatening infections associated with fungal pathogens is estimated to be on the same scale as tuberculosis and malaria with 13 million infections yearly, and a mortality of 1.5 million (Bongomin *et al*. 2017; Fisher *et al*. 2018). The severity of fungal infection is exacerbated by the widespread occurrence of immunocompromising conditions such as cancer, human immunodeficiency virus (HIV) disease, tuberculosis and the coronavirus disease 2019 (COVID-19)(Gold *et al*. 2022). Moreover, the rapid emergence of antifungal resistance in clinical strains, analogous to antibacterial resistance, has highlighted the need to expand the current toolset of treatment options targeting the main fungal pathogens, *Candida, Aspergillus* and *Cryptococcus* species. Specific strains of *Saccharomyces cerevisiae* are also opportunistic pathogens in immunocompromised patients (Perez-Torrado and Querol 2015). The impact of the rise in antifungal resistance is now recognised as an emerging threat to public health and strategies are required to establish future mechanisms of combatting the risk caused by pathogenic fungi (Fisher *et al*. 2022).

Currently, only four classes of antifungal drugs are in routine use: azoles, polyenes, pyrimidine analogues and echinocandins (Lee *et al*. 2021). Azoles, such as fluconazole and miconazole, are the most commonly prescribed for the treatment of *Candida* and *Cryptococcus* infections (Berkow and Lockhart 2017; Lee *et al*. 2021). Azoles act as strong inhibitors of lanosterol demethylase (encoded by *ERG11*) disrupting ergosterol biosynthesis, an essential component of fungal membranes, leading to the accumulation of toxic sterol intermediates (Heimark *et al*. 2002).

Amongst the pyrimidine analogues, flucytosine has been in use as an antifungal drug since 1968 (Delma *et al*. 2021) and is often used in combination with other antifungal agents to reduce occurrences of antifungal resistance (Vermes *et al*. 2000). Flucytosine is first converted *in vivo* in 5-fluorouridine by a cytosine deaminase and then phosphorylated to 5-fluoro-uridine-triophosphate (5-F-UTP) or reduced to 5-fluoro-deoxyuridine-monophosphate (5-F-dUMP). As 5-F-UTP the drug is incorporated into RNA, leading to the inhibition of protein synthesis. Instead, as 5-F-dUMP the antifungal act as an inhibitor of thymidylate synthase and, as a consequence, of purine and DNA synthesis (Campoy and Adrio 2017; Delma *et al*. 2021).

The most recent class of antifungal introduced for clinical use are echinocandins; lipopeptides which act through a non-competitive binding of the enzyme β-D-glucan synthase complex in both *Candida* an *Aspergillus spp*. Echinocandins, such as caspofungin and micafungin, inhibit the biosynthesis of β-D-glucan, a crucial component of the fungal cell wall, leading to osmotic stress and cell death (Denning 2002).

The known molecular mechanisms of fungal resistance are often linked to mutations in the target gene or in the pathway affected by the drug. As a case in point, *Candida spp*. are known to develop resistance to fluconazole by the overexpression of the target gene, *ERG11*, or by the accumulation of point mutations which alter the structure of the molecular target. However, the high-level of resistance of clinical isolates can rarely be ascribed to the effect of single mutations, which, instead, is often attributable to a combination of traits and a gradual adaptation to the stressor (Berkow and Lockhart 2017; Berkow *et al*. 2020).

A population response to an environmental stressor is impacted by the genetic variation present within the gene pool, where there exists a spectrum of fitness based on allelic determination. This is true of all organisms including pathogenic fungi and can be associated to antifungal drug susceptibility. The strong selection pressures of exposure to such agents can result in the propagation of antifungal resistance (Fisher *et al*. 2018), the emergence of fluconazole resistant *C. albicans* strains is well documented in long-term fluconazole treatment in HIV positive individuals (Johnson *et al*. 1995). Indeed, exposure to different concentrations of drugs can also influence the speed and nature of the adaptive response of pathogenic fungi and yeasts to antifungal drugs (Todd *et al*. 2023). Therefore, to perform thorough investigations of the networks and pathways associated with antifungal resistance it is essential to carry out genome-wide investigations to sample the breadth of the cellular response to antifungal agents.

In recent years, a growing number of high-throughput and -omics studies have been successfully applied to the study of antifungal resistance and pathogenicity in both model systems, such as *Saccharomyces cerevisiae* and *Schizosaccharomyces pombe* (Zhang *et al*. 2002; Ehrenreich *et al*. 2010; Zhou *et al*. 2013; Phadke *et al*. 2018), and pathogenic species such as *C. albicans* and *Cryptococcus neoformans* (Homann *et al*. 2009; Vogan *et al*. 2016). However, such studies are often limited by the use of laboratory strains which precludes in-depth analysis into the mechanisms developing through adaptation and evolution in nature. To establish new routes of therapeutic treatment we require methods to identify genetic targets for antifungal drugs that consider the variation present across different species and natural populations of Ascomycota.

Here, we apply a pipeline for the identification of QTLs in multigenerational yeast hybrids to unpick allelic variants associated with antifungal susceptibility. Diploid hybrid spores derived from crossing of strains belonging to two different species, namely *S. cerevisiae* and *S. kudriavzevii*, which were propagated for 12 meiotic generations (F12; (N_ASEEB_ *et al*. 2021), were tested for growth on antifungals. The offspring revealed a broad phenotypic diversity in the majority of conditions tested, highlighting the large impact that gene interactions have on fitness. The QTL study identified hybrid-specific traits linked to resistant phenotype in fluconazole, flucytosine and micafungin, highlighting how natural variations and hybridization could lead to resistant phenotypes. We found genes that are drug specific and two pleiotropic genes, *BCK2* and *DNF1*, that affect the phenotype across all three drugs. Moreover, we identify 41 QTL regions that contain genes not currently associated with drug resistance in the published literature. The data generated represent a valuable resource for the identification of new markers and predictors of antifungal resistance and to inform drug development.

## Material and methods

### Strains used in the study

The parental tetraploid *S. cerevisiae* x *S. kudriavzevii* hybrid (Sc^OS253^/Sk^OS575^/Sc^OS104^/Sk^IF01802^) harbouring *S. kudriavzevii* mitochondria and 228 F12 diploid hybrid progeny generated in Naseeb et al. 2021 were used in this study. Yeast strains were maintained in YPD medium (1% yeast extract, 2% peptone, 2% glucose, Formedium, UK) in 96 well-plates, and in PlusPlates (Singer Instruments, UK) with YPD + 2% agar incubated at 30°C.

### Phenotypic tests

Diploid spores of *S. cerevisiae* x *S. kudriavzevii* hybrids were grown in YPDA at 30°C and then inoculated into 100 µL YPD in a 96 well microtiter plate alongside the parental tetraploid (Sc^OS253^/Sk^OS575^/Sc^OS104^/Sk^IF01802^). The strains were incubated for 96 hours at 30°C and then sub-cultured in 384 well microtiter plate containing 70 µL YPD using the Singer Rotor HAD (Signer Instruments, Somerset, UK), prepared with four technical replicates of each strain.

For the fitness analysis of *S. cerevisiae* x *S. kudriavzevii* spores, the liquid cultures were grown to saturation at 30°C and stamped in solid media plates at a density of 384 strains per plate. The spores were grown at 30°C in: YPDA, YPDA + 5 μg/ml and 10 μg/ml of fluconazole, 10 μM and 20 μM of miconazole, 1 mg/ml and 2.5 mg/ml of caffeine, 25 ng/ml and 50 ng/ml of micafungin, 10 μg/ml and 20 μg/ml of flucytosine, 0.5 μg/ml and 1 μg/ml of bleomycin.

The plates were imaged with the PhenoBooth Colony Counter (Singer Instruments, UK) after 72 hours of incubation and the size of the individual colonies was used as a proxy for fitness. DNA Extraction and sequencing

Diploid spores of *S. cerevisiae* x *S. kudriavzevii* hybrids were inoculated in 1.5 mL of YPD and incubated overnight at 30°C, shaking. Total DNA was purified using Epicentre Masterpure™ Yeast DNA Purification Kit (Lucigen, USA) and resuspended in 50 µL of RNase-free water. To remove any RNA contamination present in the sample, the purified DNA was incubated for 30 minutes at 37°C with 1 µL of 5 µg/µL RNase A.

The quality of the purified DNA was assessed through gel electrophoresis on a 0.8% agarose gel and a Thermo Scientific™ NanoDrop Lite spectrophotometer (Thermo Scientific, UK). The DNA in each sample was quantified with a Qubit 4 Fluorometer (Thermo Scientific, UK).

### QTL mapping

The unmapped paired-end sequences from an Illumina HiSeq 4000 sequencer were quality assessed by FastQC (Andrews 2010). Sequence adapters were removed, and reads were quality trimmed (to quality score q20) using Trimmomatic_0.36 (Bolger *et al*. 2014). The mapping, variant calling and Multipool analysis was performed as previously described (Naseeb *et al*. 2021). Briefly, the reads were mapped against a reference hybrid genome containing the reference sequence for each founder species *S. cerevisiae* OS104 (Yue *et al*. 2017) and *S. kudriavzevii* IF0 1802T Ultra-Scaffolds assembly (Scannell *et al*. 2011) using BWA-MEM (Li and Durbin 2009) (bwa_ 0.7.15). Local realignment was performed with GATK_3.8.0 (DePristo *et al*. 2011) and duplicates were marked with Picard Toolkit_2.1.0 (http://broadinstitute.github.io/picard/). The alignment quality was assessed with Qualimap_2.2.1 (García-Alcalde *et al*. 2012). For each sample, variant calling was performed individually using Freebayes_1.1.0 (Garrison E 2012) with ploidy setting at 1 and including the following parameters --minmapping-quality 30 --min-base-quality 20 --no-mnps. The resultant VCF files were filtered for ‘type=SNP’ variants and processed using R. Unique bi-allele markers for each founder species were identified. Reads depths below 10 were excluded. The parental allele counts in each F12 pool were then calculated by matching reference (RO) and alternate (AO) alleles to the bi-allele marker sets among the founders.

The allele counts were provided to Multipool (Edwards and Gifford 2012) and log_10_ likelihood ratios (LOD scores) were calculated across each chromosome. QTLs were identified and reported with an LOD support interval of 1, in regions where a minimum LOD score of 5 extended across at least 20kb. The identified QTLs were annotated using the available annotation from *S. cerevisiae* OS104. Annotation for the *S. kudriavzevii* IF0 1810T ultra-scaffolds assembly was performed using HybridMine (Timouma *et al*. 2020). The pipeline for QTL analysis can be found at https://github.com/Sookie-S/QTL_analysis_pipeline_hybrid_species/tree/main.

### Data analysis

Potential causal genes were analysed with the Sorting Intolerant from Tolerant (SIFT) algorithm to assess if amino-acids variants were predicted to influence the protein function. SIFT analysis were conducted using data from Bergstrom et al. 2014 (Bergstrom *et al*. 2014) on the *S. cerevisiae* strains OS104 and OS253.

### Validation through RHA

Reciprocal hemizygosity analysis (RHA) provides a method of validation for identified, advantageous, alleles (Steinmetz *et al*. 2002) and was broadly performed as in Naseeb at al. 2021 (Naseeb *et al*. 2021). RHA was performed on candidate genes *BCK2* and *DNF1*. PCR-mediated deletion of the *S. cerevisiae* allele was performed in the F12 *Saccharomyces cerevisiae*/*Saccharomyces kudriavzevii* diploid hybrids (Sc^OS104^/Sk^IF01802^ and Sc^OS253^/Sk^OS575^) and the *S. cerevisiae* strains (Sc^OS104^ and Sc^OS253^). All deletion strains were confirmed using confirmation colony PCR (all primers used are included in table S6). Mass mating was applied to generate reciprocal hemizygotes for both interspecies tetraploids and intraspecies diploids, hybrids were selected for on triple drug plates (300 μg/mL geneticin, 200 μg/mL nourseothricin and 15 μg/mL phleomycin).

The fitness of *DNF1* and *BCK2* allele variants was tested in liquid YPD media with added flucytosine (50 μg/mL) and micafungin (50 ng/mL) on a FLUOstar Optima 466 plate reader (BMG) at 25°C. The growth characteristics of the plate reader experiments were assessed with the R package Growthcurver using K as maximum biomass, r as maximum growth rate, auc_l as integral area and Tmid as the time at which the population density reaches 1/K. Statistics were performed using GraphPad Prism.

## Results and Discussion

### Phenotypic screening of inter-specific hybrid offspring in the presence of antifungal drugs

Multigenerational diploid offspring of tetraploid hybrids (Naseeb *et al*. 2021) containing the genome of four strains belonging to two different species (Table 1), namely *S. cerevisiae* strain OS104 (Sc^OS104^) and OS253 (Sc^OS253^), and *S. kudriavzevii* strain IFO1802 (Sk^IF01802^) and OS575 (Sk^OS575^) were used to investigate growth changes in the presence of antifungals. High-throughput phenotyping on solid media (Barton *et al*. 2018) allowed us to assess how genotypic divergence of the hybrid spores impacts resistance or susceptibility to antifungal agents. The growth of the hybrid progenies along with their tetraploid parent was assessed in standard rich medium and in media containing inhibitors of ergosterol biosynthesis (fluconazole and miconazole), cell wall biogenesis (micafungin and caffeine) and nucleic acid biosynthesis (flucytosine and phleomycin). The antifungal drugs were added at sub-lethal concentrations in the media to allow for two-tailed selection to identify both high-performing and low-performing spores.

**Table 1.** Background and origins of all parental strains and hybrid tetraploids used for phenotypic screening.

| Species and hybrids | Strain | Origin |
| --- | --- | --- |
| *Saccharomyces cerevisiae* | Sc<sup>OS253</sup> (NRRL-Y12663) | Sake (Palm wine, Africa) |
|  | Sc<sup>OS104</sup> (YPS128) | North American (woodland isolate) |
| *Saccharomyces kudriavzevii* | Sk<sup>OS575</sup> (48BYC-4) | China, oak bark |
|  | Sk<sup>IFO1802</sup> (NCYC2889) | Japan, Decayed leaf matter |
| Sc/Sk Tetraploid hybrid | Sc<sup>OS253</sup>/Sk<sup>OS575</sup>/Sc<sup>OS104</sup>/Sk<sup>IFO1802</sup> | DD's lab |

The hybrid progeny exhibited a broad phenotypic space in all the six antifungal drugs screened (Figure 1, Table S1, Figure S2). The higher dispersion was recorded in media with 10 μg/ml of fluconazole with a IQR (quartile coefficient of dispersion) of 0.55, compared to 0.14 in YPD (Table S1). The growth in miconazole and phleomycin, at 1μg/mL, were characterised by low cell viability (<99%), with only around 20% of the progeny able to sustain growth at the highest concentration tested (Table S1). The tetraploid parent control was able to grow in each of the antifungal drugs tested except for miconazole. Compared to its progeny, it performed worse than the than the median of the offspring in four out of six conditions (present in the upper quartile only in phleomycin and fluconazole; Figure 1). These data highlight the transgressive traits of the hybrid offspring compared to the parent, and support the notion that the greater fitness of progeny may be the result of unlinking detrimental alleles present in the parental tetraploid.

**Figure 1.**
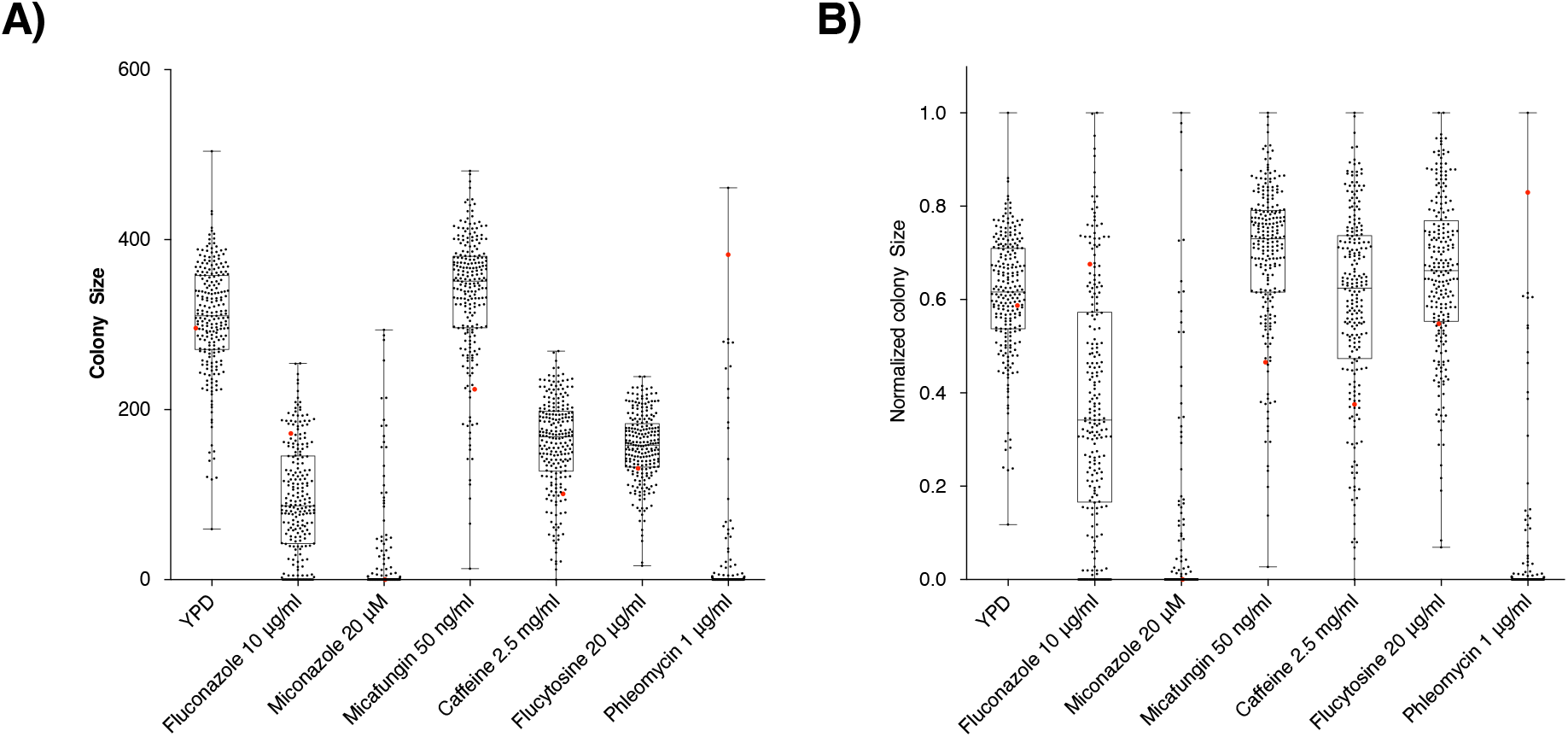
Box plot of the fitness of F12 diploid progeny for *S. cerevisiae/S. kudriavzevii*. hybrids after incubation with different concentrations of antifungal drugs expressed as colony size (A) and as normalized colony size (B) parental tetraploid was used as a control and is highlighted in red. Each black dot represents a distinct F12 hybrid progeny. The upper and lower bars show the maximum and minimum values, the box represents the second and third quartile with the central line the median.

The hybrid spores grown in media containing micafungin exhibited a higher median colony size compared to growth in YPD (Figure 1A, Table S1, Figure S2). Previous studies observed an abnormal morphology in yeast cells treated with micafungin, reportedly similar to *HOC1* and *MNM10* mutants, which correlated with an abnormally large morphology (Jorgensen *et al*. 2002; Gebre *et al*. 2015). Therefore, the larger colony size observed may be correlated to an increase in cell size within the population.

### Identification of QTL regions associated with susceptibility to antifungal agents

To identify the genetic traits underlying antifungal resistance in the hybrid progeny, we used the multipool approach (Cubillos *et al*. 2013) and sequenced a pool of the 20 best and of the 20 worst performing strains for three antifungal drugs acting on different biological processes, namely fluconazole, micafungin and flucytosine. The resulting genomic data was analysed following an in-house pipeline (Figure S1, Github) which allowed the identification of QTL regions where allele frequencies significantly differed between the two pools of spores and exhibited a diametrically opposite trend.

Across the three drugs, a total of 101 and 89 different QTL regions were identified in the *S. cerevisiae* and in the *S. kudriavzevii* genomes, respectively (Table 2). These results reinforce the notion that while usually antifungals target a specific enzyme or metabolic process, the genetic background has a strong influence on the degree of drug resistance/susceptibility shown by the host. The largest number of QTLs was detected in flucytosine with 47 and 51 QTL regions identified in *S. cerevisiae* and *S. kudriavzevii*, respectively. The higher number of QTL associated with the trait may result from a higher complexity of the target of flucytosine, which is known to affect both RNA and DNA biosynthesis in fungi (Vermes *et al*. 2000).

**Table 2:** Number of QTL regions and genes identified in S. *cerevisiae* (Sc) and *S. kudriavzevii* (Sk) grown on different drugs.

| Genome | Number of QTL regions detected |  | Number of genes within the QTL regions |  |
| --- | --- | --- | --- | --- |
|  | Sc | Sk | Sc | Sk |
| **Micafungin** | 25 | 17 | 93 | 48 |
| **Fluconazole** | 29 | 21 | 212 | 107 |
| **Flucytosine** | 47 | 51 | 175 | 279 |
| **Total** | 101 | 89 | 480 | 434 |

SIFT (Sorting Intolerant from Tolerant) algorithm was used to identify non-synonymous SNPs between the two parental alleles of *S. cerevisiae* that could affect the protein function (Kumar *et al*. 2009; Bergstrom *et al*. 2014). Around 56% of genes identified within the QTL regions had one or more SNPs between the parental strains (Table S2).

### Identification of potential causal genes within QTL regions

A total of 136 genes (Table 3) have been identified in QTL regions that have potential to affect the phenotype based on previously published literature. Of the 136 genes, 75 were identified across QTL regions within either *Sc* and *Sk* genomes that have previously found to have a direct role in resistance to antifungal drugs (fluconazole, flucytosine, phleomycin, caspofungin and echinocandin) based on classical genetic (Cherry *et al*. 2012) or transcriptome studies (Zhang *et al*. 2002) in *S. cerevisiae* (Table 2). Fifty-six were found interacting with the drug target (*e*.*g*. with *ERG11* in fluconazole-QTLs) or with cellular processes closely linked to the drug mechanism of action (*e*.*g*. nucleotide biosynthetic processes for flucytosine-QTLs or β-glucan metabolism for micafungin-QTLs). Moreover, in micafungin-QTLs, we identified four orthologues of *Schizosaccharomyces pombe* genes associated with micafungin resistance (Zhou *et al*. 2013). Amongst these, *TOR1*, a protein kinase involved in signal transduction, cell growth and autophagy, was found similarly involved in the development of resistance to caffeine, a cell wall stressor, in both *S. cerevisiae* and *C. albicans* (Homann *et al*. 2009).

**Table 3:** List of 136 causal genes identified in QTL regions. Genes are clustered depending on their role or by the phenotype recorded in genetic studies. 75 genes are listed for their known role in resistance to antifungal drugs, further genes are listed for their role in pathways associated with antifungal drug susceptility, including 30 genes with physical or genetic interactions with *ERG11*.

| Causal genes identified in Fluconazole QTLs |  |  |
| --- | --- | --- |
|  | *S. cerevisiae* alleles | *S. kudriavzevii* alleles |
| Increased resistance to fluconazole | *PPZ1, SLY41, YAP1, ARP8, DEG1, HST4, ERV25, RAD17, ISW2, HAP5, ERG6, MRT4, SNU66, ADR1, GYP1, DST1, PSP2, MSA1, ANY1, NUP42, MRX16, TAF12, UBP6, CDC5, ITT1, UPC2, ALG8, PDE2, NDD1, COM2, HSC82, SWI5, COQ4, LDB19, VTS1, PRT1* | *SSK1, PUB1, RDH54, KES1, VPS27, CSE2, ACS2, ARK1, CBF2, HDA1, SOL1* |
| Ergosterol biometabolism | *ERG6, HEM2, UPC2* | *ARE2* |
| Reported physical interaction with *ERG11* | *PPZ1, ERV25, ERG6, LAC1* | *ASI3, KES1* |
| Reported genetic interaction with *ERG11* | *DOP1, TAF12, CBS2, SPC19, RAV2, RAD9, UBC6, RNA15, UBX2, SGS1, RPL36A, GYP1, HEM4, PLP2, DGK1, FAA1, PDE2, MRS6* | *NOC3, RPS10B, PET8, ASI3, LRO1, DBP6, PRP46* |
| Causal genes identified in Flucytosine QTLs |  |  |
|  | *S. cerevisiae* alleles | *S. kudriavzevii* alleles |
| Resistance to flucytosine | *BCH2, LSM6, ATO3, NKP1, ATP17, XRS2* | *PEX8, RRT2, BUB3, ROM1, NOC2, STP3, BFR1, MAK21, RRP46, PAC10, HMS1* |
| DNA Damage Repair | *RDH54, ESC2, POL1, RAD3, MLH1* | *DDR48, EXO1, IES4, MLH1* |
| Resistance to phleomycin | *CPR5, DNF1, HXK1, HNT3, BCK2, ENV11* | *SCL1, AIM34* |
| Drug resistance | *PDR15, RDS2* | *PDR10* |
| Nucleotide biosynthetic processes | *ADE8, ADK2* | *DUT1, ADE16, CDD1* |
| DE genes and paralogs after flucytosine treatment in *S. cerevisiae* | *RPL12B, ADE8, PDR15* | *MMT1, SPT21, CTL1, SER1, RPL9B, RPL15A* |
| Causal genes identified in Micafungin QTLs |  |  |
|  | *S. cerevisiae* alleles | *S. kudriavzevii* alleles |
| Resistance to caspofungin | *AKR1, MEH1* | *FEN2, MSN5, YSP2* |
| Resistance to echinocandin | *BCK2* |  |
| Beta glucan biometabolism | *EXG2* |  |
| Cell wall stress | *BCK2* | *RQC1, CNB1* |
| Resistance to caffeine | *IPK1, BCK2, TOR1* | *DPH2, DCG1, CNB1* |
| Orthologues of *S. pombe* genes associated with micafungin resistance | *MKC7, TOR1, HEL1, YTA6* |  |

Amongst potential causal genes identified from *S. cerevisiae*, SIFT analysis identified 11 alleles that carry SNPs predicted to cause a strong deleterious effect on the protein function (13.25%) while 38 were inferred to have tolerated mutations (45.8%) (Table S3). A number of QTL regions encompassed genes that have not previously been associated with drug resistance or susceptibility. These can be genes that have an impact in the original strains/species and were not previously identified, or they could be more specifically associated with the hybrid background. In total we found 41 QTL regions containing genes not previously reported in the literature for both *S. cerevisiae* and *S. kudriavzevii* (Table S4, Table S5). A total of 23 genes were identified in high LOD intervals (*i*.*e*. >16.0; Table 3), of which 11 were flagged in the SIFT analysis as harbouring non-synonymous SNPs between the parental alleles and 3 harbouring potentially deleterious effects on protein function.

### Species overlap of QTL regions found only in flucytosine-QTLs

Four QTL regions were identified in both *S. cerevisiae* and *S. kudriavzevii* alleles as affecting the hybrid fitness in flucytosine (Figure 2), while no common regions were detected in fluconazole and micafungin-QTLs. Amongst the genes mapped in the shared flucytosine-QTLs we were able to identify potential causal genes (Table 2), as their function was linked to DNA synthesis and maintenance or to drug sensitivity in previously published work. In particular, *ENV11* and *TRR1* deletions were found to affect sensitivity to phleomycin (Kapitzky *et al*. 2010; Krol *et al*. 2015), while *ALD2* null mutants had increased sensitivity to floxuridine, a pyrimidine analogue which inhibits DNA and RNA synthesis similarly to flucytosine (Lum *et al*. 2004). Moreover, *MLH1* is an ATPase involved in meiotic mismatch repair in mitosis and meiosis (Kramer *et al*. 1989) while *PFU1* overexpression was found to exacerbate UV radiation toxicity in overexpression studies (Chakrabortee *et al*. 2016).

**Figure 2.**
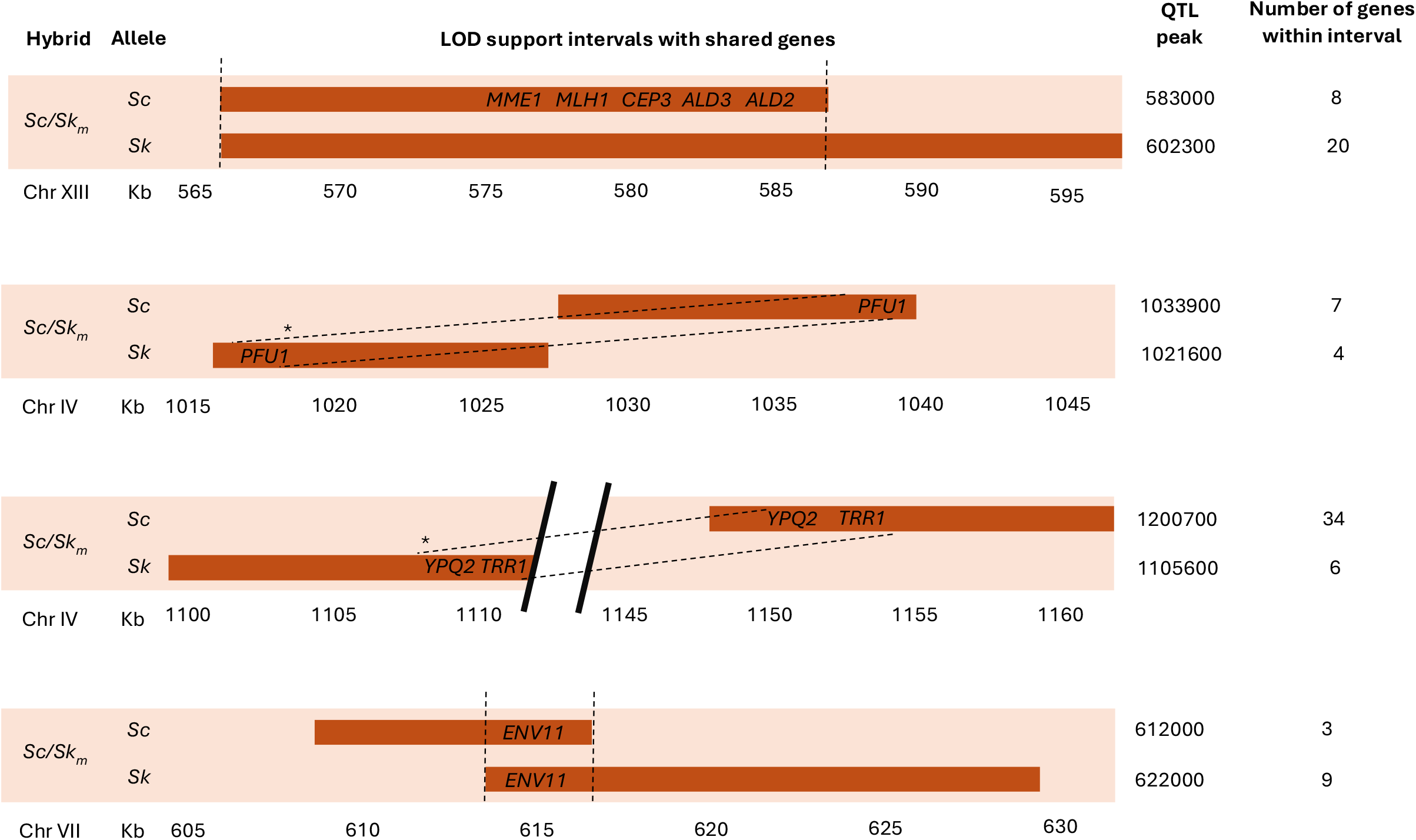
QTL intervals identified as significant for flucytosine susceptibility that are shared between *S. cerevisiae* and *S. kudriavzevii* genomes. For each interval shared, the position and the peak of the individual QTL is specified. Genes identified that are shared between *S. cerevisiae* and *S. kudriavzevii* are listed. ^*^Regions containing shared genes in chromosome IV occupy different relative positions on the chromosomes of the two yeast species.

### Pleiotropic QTLs are rare across antifungal drugs

The difference in the mechanism of action of the fluconazole, micafungin and flucytosine resulted in a high number of trait-specific QTLs (99%), with only three pleiotropic regions, involving *S. cerevisiae* alleles, shared across conditions (Table 5). A small region in the chromosome V of *S. cerevisiae*, encoding for Pab1p, Bck2p and Dnf1p, was mapped in all three conditions. *BCK2* encodes a protein involved in the regulation of the cell cycle, and was previously shown to be important for growth in the presence of cell wall stressors (*i*.*e*. caffeine and echinocandin) and antifungal drugs, such as phleomycin, affecting DNA and RNA synthesis (Kapitzky *et al*. 2010). Here, we show that *BCK2* alleles are important for growth on three further drugs, supporting the pleiotropic nature of this gene.

**Table 4.**
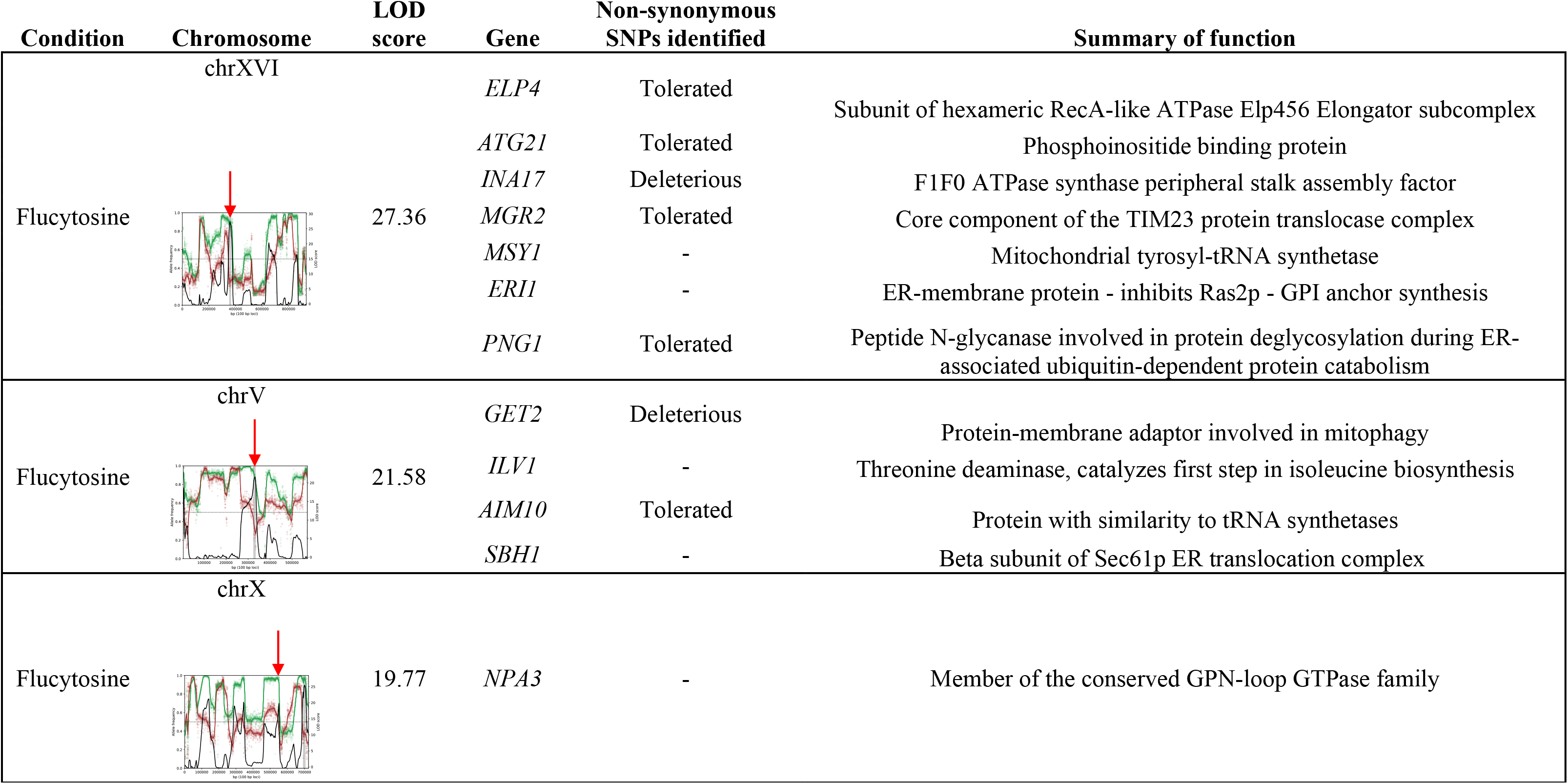
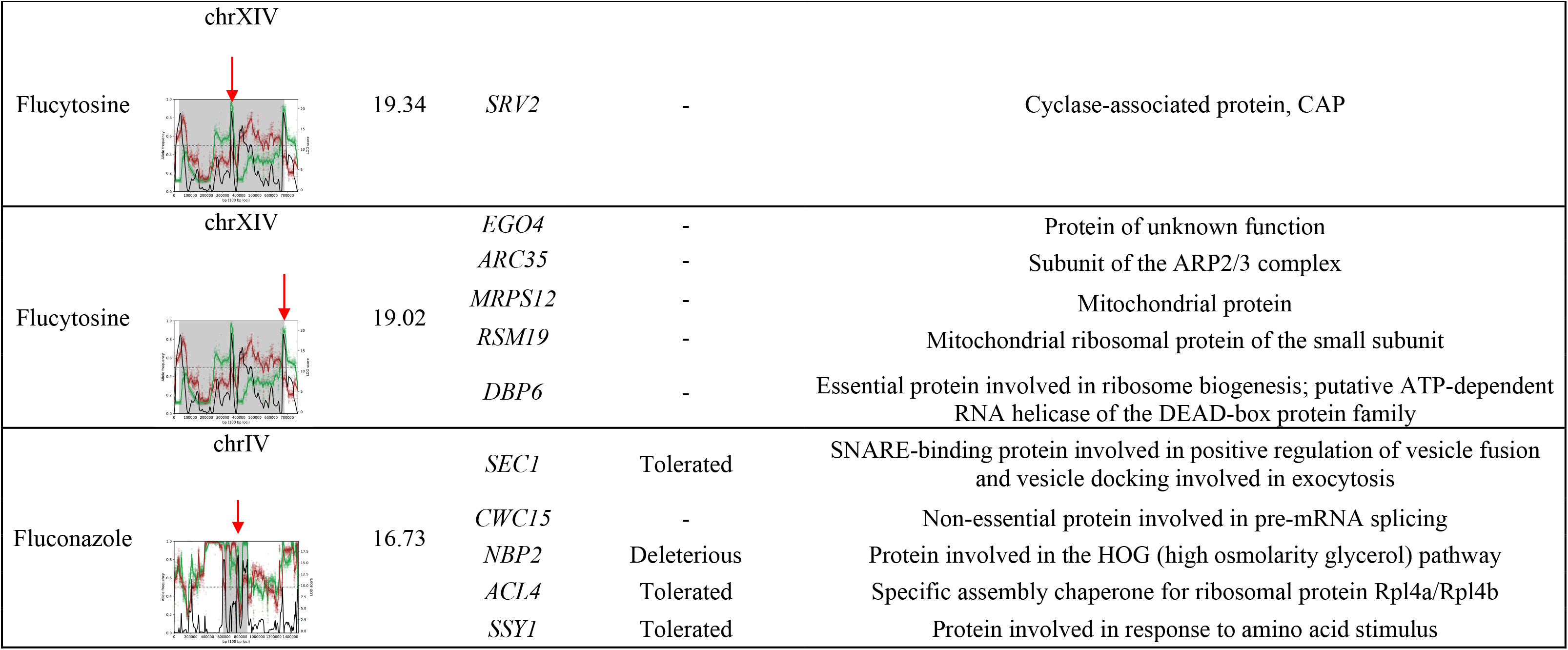
A summary of high LOD scoring intervals and the genes identified within the QTLs. Respective LOD scores, gene names and function summaries are listed for each of the regions.

**Table 5:** List of pleiotropic QTLs in *S. cerevisiae* x *S. kudriavzevii* hybrid diploid progeny. The coordinates of the pleiotropic intervals are specified along with the genome and the selection condition in which they were identified.

|  | Chromosome | Start (bp) | End (bp) | Selection conditions | Genes in QTL interval |
| --- | --- | --- | --- | --- | --- |
| *S. cerevisiae* genome | IV | 730300 | 740100 | Fluconazole<br>Micafungin | *MKC7, TAF12, SWI5, EKI1* |
|  | V | 506200 | 519700 | Fluconazole<br>Flucytosine<br>Micafungin | *PAB1, DNF1, BCK2* |
|  | XV | 592700 | 602600 | Fluconazole<br>Flucytosine | *LSC1, THI80, ELG1, PNO1* |

*DNF1*, is a flippase involved in the phospholipid translocation, and its deletion increase the fitness of *S. cerevisiae* grown on phleomycin (Kapitzky *et al*. 2010). Additionally, it has been shown that the deletion of *DNF1* orthologue in *Nakaseomyces glabratus* (formerly *Candida glabrata*) confer resistance to caspofungin (Rosenwald *et al*. 2016). Again, our data also highlight the pleiotropic nature of *DNF1*. Interestingly, both *DNF1* and *BCK2* are linked and have a role in the antifungal resistance trait, such linking of alleles in yeast may be an adaption to limit the segregation of advantageous allele combinations during recombination analogous to ‘supergenes’ in butterflies (Liti and Louis 2012).

### Validation of QTLs by Reciprocal hemizygosity analysis

We chose to validate two genes that were identified for their pleiotropic role, namely *BCK2* and *DNF1*, whose alleles present tolerated and deleterious non-synonymous SNPs, respectively, according to the SIFT analysis. Dnf1p is a flippase involved in phospholipid translocation, and Bck2p is a protein involved in the G1/S transition of the cell cycle (Wijnen and Futcher 1999). *DNF1* and *BCK2* were validated by reciprocal hemizygosity analysis (RHA; Fig. 3A) in flucytosine and micafungin, respectively, based on the LOD score with the highest for the respective QTLs in these two conditions. We created reciprocal allelic deletions of *DNF1* and *BCK2* (*i*.*e*. Sc^D^/Sk^IF01802^/Sc^OS253^/Sk^OS575^; Sc^OS104^/Sk^IF01802^/Sc^D^/Sk^OS575^) in the tetraploid parent (Sc^OS104^/Sk^IF01802^/Sc^OS253^/Sk^OS575^) and compared the growth of mutants in the presence of antifungal (Figure 3B).

**Figure 3.**
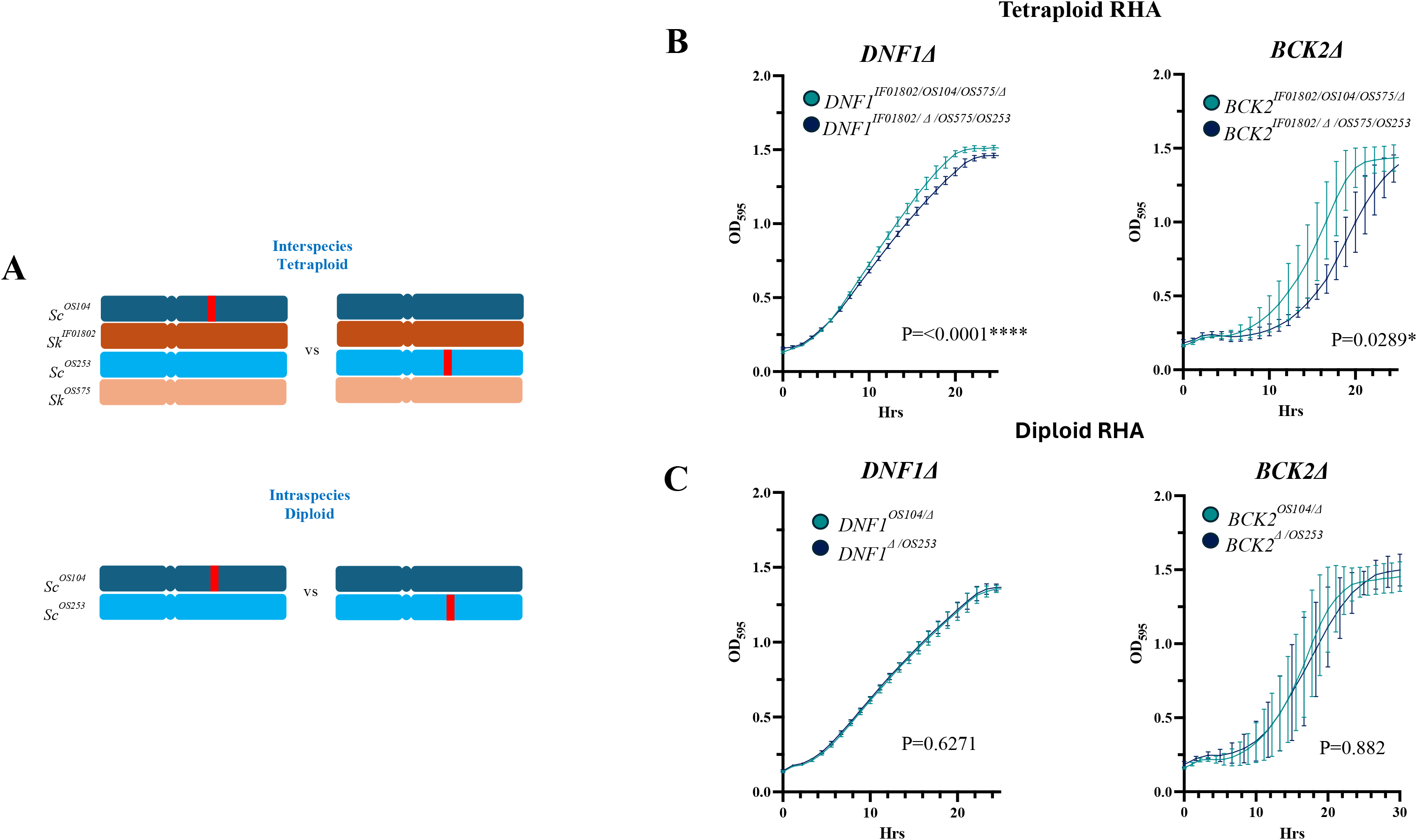
Growth curves for inter- and intra-specific hemizygotes assessed in antifungal drugs. **A** Diagram representing the process of reciprocal hemizygote generation for interspecies tetraploids, the GOI is knocked out via PCR-mediated deletion in the interspecies diploid. Mating with the WT diploid results in a hemizygote tetraploid for the Sc alleles. **B** Growth curves for Δ*DNF1* ScSk_m_ tetraploid reciprocal hemizygotes and Δ*BCK2* ScSk_m_ tetraploid reciprocal hemizygotes **C** Growth curves for the Δ*DNF1* ScSc diploids and the growth curve for the Δ*BCK2* ScSc diploids is shown. Fitness assays were performed in YPD media + 50ug/mL flucytosine (*DNF1*) or YPD + 50ng/mL micafungin (*BCK2*) as highlighted in materials and methods. The integral area of the growth curves was calculated using the Growthcurver R package and significance assessed by the Student’s t-test (P>0.05).

We observed a significant difference in maximum growth rate (*P*=0.0001^***^) and final biomass (P=0.0001) between *DNF1* alleles (Figure 3B, Table S7), and a significant difference in the lag phase for the *BCK2* alleles (Figure 3B). In both cases, there was a significant difference of the integral area under the growth curve (Figure 3B, Table S7): the *DNF1*^OS104^ allele performed significantly better than the *DNF1*^OS253^ allele in the presence of flucytosine (Figure 3B, Table S7; *P*=0.0001^***^), reflecting the QTL genetic screening; the *BCK2*^OS104^ allele performed better than *BCK2*^OS253^ in response to micafungin (Figure 3B; *P*=0.0289^*^).

Here the direction of the fitness was opposite to that observed in the QTL data, this is not a concerning observation as RHA was performed on the hemizygote tetraploids and phenotypic screening performed on diploid progeny. Genetic interactions in tetraploids differ to the diploid progeny, differences in copy number and negative and positive epistasis can influence the phenotype (Greig *et al*. 2002). Such transgressive segregation is widely observed in plants, where progeny show both fitter characteristics and less fit phenotypes than the parent (Rieseberg *et al*. 1999). The F12 tetraploids may demonstrate an altered phenotype in comparison to the diploid progeny as all alleles, both advantageous and deleterious, are present within one strain.

### Antifungal QTLs are specific to interspecies hybrids

To establish whether the QTLs identified are specific to the interspecies hybrid background, we tested the *BCK2* and *DNF1* allelic variants in two strains of the same species. We performed RHA using the parental *S. cerevisiae* strains *Sc*^*OS104*^ and *Sc*^*OS253*^, generating hemizygotic diploids (*Sc*^*OS104*^*/Sc*^D^ and *Sc*^D^/*Sc*^*OS253*^) with deletions of *BCK2* and *DNF1* alleles (Fig. 3 A). Between allelic variants, we observed no significant difference in growth across both conditions (Figure 3C, Table S8) with no significant difference in the integral area under the growth curve in both conditions (P>0.05), or for final biomass (Figure 3C, Table S8). Through RHA, we were able to validate the phenotypic effect of the candidate genes and identify allelic variants that represent markers of antifungal resistance. As previously noted in Naseeb et al. 2021 (Naseeb *et al*. 2021), new and unique QTLs are identified in hybrids, we have confirmed that two genes identified in interspecies progeny do not have the same phenotypic effect in the *Sc/Sc* diploid background. Hybrid models, such as that developed by Timouma *et al*. for the hybrid *Saccharomyces pastorianus* (Soukaina Timouma 2023), are needed to understand the emergence of allelic variants that impact phenotype exclusively in hybrids.

## Conclusions

This study allowed us to investigate how natural variation, developing outside of the clinical samples, may affect the evolutionary pathway leading to a resistant phenotype. We exploited recent advances multigenerational breeding of *Saccharomyces* interspecies hybrids to study the impact of the allelic variants on the resistance and susceptibility to a diverse range of antifungal drugs, including azoles, pyrimidine analogues and echinocandins. We demonstrated the broad phenotypic diversity of the diploid hybrid progeny across the conditions tested and have developed a robust pipeline for the identification of QTL regions in yeast interspecies hybrids. We identified major QTLs for the different drugs, including some not previously associated with antifungal resistance. Three pleiotropic regions, involving *S. cerevisiae* alleles, were shared across all drug conditions. We validated, via reciprocal hemizygosity analysis, the impact of allele variants of two pleiotropic genes, namely *BCK2* and *DNF1*, in response to micafungin and flucytosine, respectively, and showed that the phenotype was hybrid specific. Interestingly, these two genes are linked suggesting an evolutionary trajectory to limit the independent segregation of advantageous allele combinations (Liti and Louis 2012).

Understanding the unique genetic interactions underlying the phenotype observed in hybrid organisms will have an impact on both drug development and personalised treatments. Furthermore, the dissection of the pleiotropic QTLs could allow the identification of new strong predictors of drug resistance. The outlined approach will be also useful to tackle human pathogens, such as *Candida* sp. and *Cryptococcus* sp., where hybridisation is commonplace (Samarasinghe *et al*. 2020).

## Data Availability

Sequencing data have been deposited in the European Nucleotide Archive (https://www.ebi.ac.uk/ena/browser/view/PRJEB70903). All other study data are included in the article and/or supporting information.

## Acknowledgments

This work was supported by the Future Biomanufacturing Research Hub (Future BRH), funded by the Engineering and Physical Sciences Research Council (EPSRC) and Biotechnology and Biological Sciences Research Council (BBSRC) as part of UK Research and Innovation (grant EP/S01778X/1); and by internal funding for Research Technology & Development awarded to DD and LZ. FV and ST were supported by H2020-MSCA-ITN-2017 grant to DD (764364; https://cordis.europa.eu/project/id/764364). WR is supported by a studentship from the Future BRH and Singer Instruments. We wish to thank the Bioinformatics Core Facility at the University of Manchester for the sequencing.

## References

Andrews, S., 2010 FastQC A Quality Control Tool for High Throughput Sequence Data.

Barton, D. B. H., D. Georghiou, N. Dave, M. Alghamdi, T. A. Walsh et al., 2018 PHENOS: a high-throughput and flexible tool for microorganism growth phenotyping on solid media. BMC Microbiol 18: 9.

Bergstrom, A., J. T. Simpson, F. Salinas, B. Barre, L. Parts et al., 2014 A high-definition view of functional genetic variation from natural yeast genomes. Mol Biol Evol 31: 872–888.

Berkow, E. L., and S. R. Lockhart, 2017 Fluconazole resistance in Candida species: a current perspective. Infection and Drug Resistance 10: 237–245.

Berkow, E. L., S. R. Lockhart and L. Ostrosky-Zeichner, 2020 Antifungal Susceptibility Testing: Current Approaches. Clinical Microbiology Reviews 33: e00069–00019.

Bolger, A. M., M. Lohse and B. Usadel, 2014 Trimmomatic: a flexible trimmer for Illumina sequence data. Bioinformatics (Oxford, England) 30: 2114–2120.

Bongomin, F., S. Gago, R. O. Oladele and D. W. Denning, 2017 Global and Multi-National Prevalence of Fungal Diseases—Estimate Precision. Journal of Fungi 3: 57.

Campoy, S., and J. L. Adrio, 2017 Antifungals. Biochemical Pharmacology 133: 86–96.

Chakrabortee, S., J. S. Byers, S. Jones, D. M. Garcia, B. Bhullar et al., 2016 Intrinsically Disordered Proteins Drive Emergence and Inheritance of Biological Traits. Cell 167: 369-381.e312.

Cherry, J. M., E. L. Hong, C. Amundsen, R. Balakrishnan, G. Binkley et al., 2012 Saccharomyces Genome Database: the genomics resource of budding yeast. Nucleic Acids Research 40: D700–705.

Cubillos, F. A., L. Parts, F. Salinas, A. Bergstrom, E. Scovacricchi et al., 2013 High-resolution mapping of complex traits with a four-parent advanced intercross yeast population. Genetics 195: 1141–1155.

Delma, F. Z., A. M. S. Al-Hatmi, R. J. M. Brüggemann, W. J. G. Melchers, S. de Hoog et al., 2021 Molecular Mechanisms of 5-Fluorocytosine Resistance in Yeasts and Filamentous Fungi. Journal of Fungi 7: 909.

Denning, D. W., 2002 Echinocandins: a new class of antifungal. Journal of Antimicrobial Chemotherapy 49: 889–891.

DePristo, M. A., E. Banks, R. Poplin, K. V. Garimella, J. R. Maguire et al., 2011 A framework for variation discovery and genotyping using next-generation DNA sequencing data. Nature Genetics 43: 491–498.

Edwards, M. D., and D. K. Gifford, 2012 High-resolution genetic mapping with pooled sequencing. BMC Bioinformatics 13 Suppl 6: S8.

Ehrenreich, I. M., N. Torabi, Y. Jia, J. Kent, S. Martis et al., 2010 Dissection of genetically complex traits with extremely large pools of yeast segregants. Nature 464: 1039–1042.

Fisher, M. C., A. Alastruey-Izquierdo, J. Berman, T. Bicanic, E. M. Bignell et al., 2022 Tackling the emerging threat of antifungal resistance to human health. Nat Rev Microbiol 20: 557–571.

Fisher, M. C., N. J. Hawkins, D. Sanglard and S. J. Gurr, 2018 Worldwide emergence of resistance to antifungal drugs challenges human health and food security. Science 360: 739–742.

García-Alcalde, F., K. Okonechnikov, J. Carbonell, L. M. Cruz, S. Götz et al., 2012 Qualimap: evaluating next-generation sequencing alignment data. Bioinformatics (Oxford, England) 28: 2678–2679.

Garrison E. M. G., 2012 Haplotype-based variant detection from short-read sequencing. arXiv 2012;1207.3907.

Gebre, A. A., H. Okada, C. Kim, K. Kubo, S. Ohnuki et al., 2015 Profiling of the effects of antifungal agents on yeast cells based on morphometric analysis. FEMS Yeast Research 15: fov040.

Gold, J. A. W., F. B. Ahmad, J. A. Cisewski, L. M. Rossen, A. J. Montero et al., 2022 Increased Deaths From Fungal Infections During the Coronavirus Disease 2019 Pandemic—National Vital Statistics System, United States, January 2020–December 2021. Clinical Infectious Diseases: ciac489.

Greig, D., R. H. Borts, E. J. Louis and M. Travisano, 2002 Epistasis and hybrid sterility in Saccharomyces. Proc Biol Sci 269: 1167–1171.

Heimark, L., P. Shipkova, J. Greene, H. Munayyer, T. Yarosh-Tomaine et al., 2002 Mechanism of azole antifungal activity as determined by liquid chromatographic/mass spectrometric monitoring of ergosterol biosynthesis. Journal of mass spectrometry: JMS 37: 265–269.

Homann, O. R., J. Dea, S. M. Noble and A. D. Johnson, 2009 A Phenotypic Profile of the Candida albicans Regulatory Network. PLOS Genetics 5: e1000783.

Johnson, E. M., D. W. Warnock, J. Luker, S. R. Porter and C. Scully, 1995 Emergence of azole drug resistance in Candida species from HIV-infected patients receiving prolonged fluconazole therapy for oral candidosis. J Antimicrob Chemother 35: 103–114.

Jorgensen, P., J. L. Nishikawa, B.-J. Breitkreutz and M. Tyers, 2002 Systematic Identification of Pathways That Couple Cell Growth and Division in Yeast. Science 297: 395–400.

Kapitzky, L., P. Beltrao, T. J. berens, N. Gassner, C. Zhou et al., 2010 Cross-species chemogenomic profiling reveals evolutionarily conserved drug mode of action. Molecular Systems Biology 6: 451.

Kramer, B., W. Kramer, M. S. Williamson and S. Fogel, 1989 Heteroduplex DNA correction in Saccharomyces cerevisiae is mismatch specific and requires functional PMS genes. Molecular and Cellular Biology 9: 4432–4440.

Krol, K., I. Brozda, M. Skoneczny, M. Bretne and A. Skoneczna, 2015 A Genomic Screen Revealing the Importance of Vesicular Trafficking Pathways in Genome Maintenance and Protection against Genotoxic Stress in Diploid Saccharomyces cerevisiae Cells. PLOS ONE 10: e0120702.

Kumar, P., S. Henikoff and P. C. Ng, 2009 Predicting the effects of coding non-synonymous variants on protein function using the SIFT algorithm. Nat Protoc 4: 1073–1081.

Lee, Y., E. Puumala, N. Robbins and L. E. Cowen, 2021 Antifungal Drug Resistance: Molecular Mechanisms in Candida albicans and Beyond. Chemical reviews 121: 3390–3411.

Li, H., and R. Durbin, 2009 Fast and accurate short read alignment with Burrows-Wheeler transform. Bioinformatics 25: 1754–1760.

Liti, G., and E. J. Louis, 2012 Advances in quantitative trait analysis in yeast. PLoS Genet 8: e1002912.

Lum, P. Y., C. D. Armour, S. B. Stepaniants, G. Cavet, M. K. Wolf et al., 2004 Discovering Modes of Action for Therapeutic Compounds Using a Genome-Wide Screen of Yeast Heterozygotes. Cell 116: 121–137.

Naseeb, S., F. Visinoni, Y. Hu, A. J. H. Roberts, A. Maslowska et al., 2021 Restoring fertility in yeast hybrids: Breeding and quantitative genetics of beneficial traits. Proceedings of the National Academy of Sciences 118.

Perez-Torrado, R., and A. Querol, 2015 Opportunistic Strains of Saccharomyces cerevisiae: A Potential Risk Sold in Food Products. Front Microbiol 6: 1522.

Phadke, S. S., C. J. Maclean, S. Y. Zhao, E. A. Mueller, L. A. Michelotti et al., 2018 Genome-Wide Screen for Saccharomyces cerevisiae Genes Contributing to Opportunistic Pathogenicity in an Invertebrate Model Host. G3 Genes|Genomes|Genetics 8: 63–78.

Rieseberg, L. H., M. A. Archer and R. K. Wayne, 1999 Transgressive segregation, adaptation and speciation. Heredity (Edinb) 83 (Pt 4): 363–372.

Rosenwald, A. G., G. Arora, R. Ferrandino, E. L. Gerace, M. Mohammednetej et al., 2016 Identification of Genes in Candida glabrata Conferring Altered Responses to Caspofungin, a Cell Wall Synthesis Inhibitor. G3 (Bethesda) 6: 2893–2907.

Samarasinghe, H., M. You, T. S. Jenkinson, J. Xu and T. Y. James, 2020 Hybridization Facilitates Adaptive Evolution in Two Major Fungal Pathogens. Genes 11: 101.

Scannell, D. R., O. A. Zill, A. Rokas, C. Payen, M. J. Dunham et al., 2011 The Awesome Power of Yeast Evolutionary Genetics: New Genome Sequences and Strain Resources for the Saccharomyces sensu stricto Genus. G3: Genes|Genomes|Genetics 1: 11–25.

Soukaina Timouma, L. N. B.-C., Jean-Marc Schwartz, Daniela Delneri, 2023 Development of a genome scale metabolic model for the lager hybrid yeast S. pastorianus to understand evolution of metabolic pathways in industrial settings. bioRxiv.

Steinmetz, L. M., H. Sinha, D. R. Richards, J. I. Spiegelman, P. J. Oefner et al., 2002 Dissecting the architecture of a quantitative trait locus in yeast. Nature 416: 326–330.

Timouma, S., J.-M. Schwartz and D. Delneri, 2020 HybridMine: A Pipeline for Allele Inheritance and Gene Copy Number Prediction in Hybrid Genomes and Its Application to Industrial Yeasts. Microorganisms 8: 1554.

Todd, R. T., N. Soisangwan, S. Peters, B. Kemp, T. Crooks et al., 2023 Antifungal Drug Concentration Impacts the Spectrum of Adaptive Mutations in Candida albicans. Mol Biol Evol 40.

Vermes, A., H. J. Guchelaar and J. Dankert, 2000 Flucytosine: a review of its pharmacology, clinical indications, pharmacokinetics, toxicity and drug interactions. Journal of Antimicrobial Chemotherapy 46: 171–179.

Vogan, A. A., J. Khankhet, H. Samarasinghe and J. Xu, 2016 Identification of QTLs Associated with Virulence Related Traits and Drug Resistance in Cryptococcus neoformans. G3: Genes|Genomes|Genetics 6: 2745–2759.

Wijnen, H., and B. Futcher, 1999 Genetic analysis of the shared role of CLN3 and BCK2 at the G(1)-S transition in Saccharomyces cerevisiae. Genetics 153: 1131–1143.

Yue, J.-X., J. Li, L. Aigrain, J. Hallin, K. Persson et al., 2017 Contrasting evolutionary genome dynamics between domesticated and wild yeasts. Nature Genetics 49: 913–924.

Zhang, L., Y. Zhang, Y. Zhou, Y. Zhao, Y. Zhou et al., 2002 Expression profiling of the response of Saccharomyces cerevisiae to 5-fluorocytosine using a DNA microarray. International Journal of Antimicrobial Agents 20: 444–450.

Zhou, X., Y. Ma, Y. Fang, W. Gerile, W. Jaiseng et al., 2013 A Genome-Wide Screening of Potential Target Genes to Enhance the Antifungal Activity of Micafungin in Schizosaccharomyces pombe. PLOS ONE 8: e65904.

