## Supplementary Information for "Impact of inter-species hybridisation on antifungal drug response in the *Saccharomyces* genus"

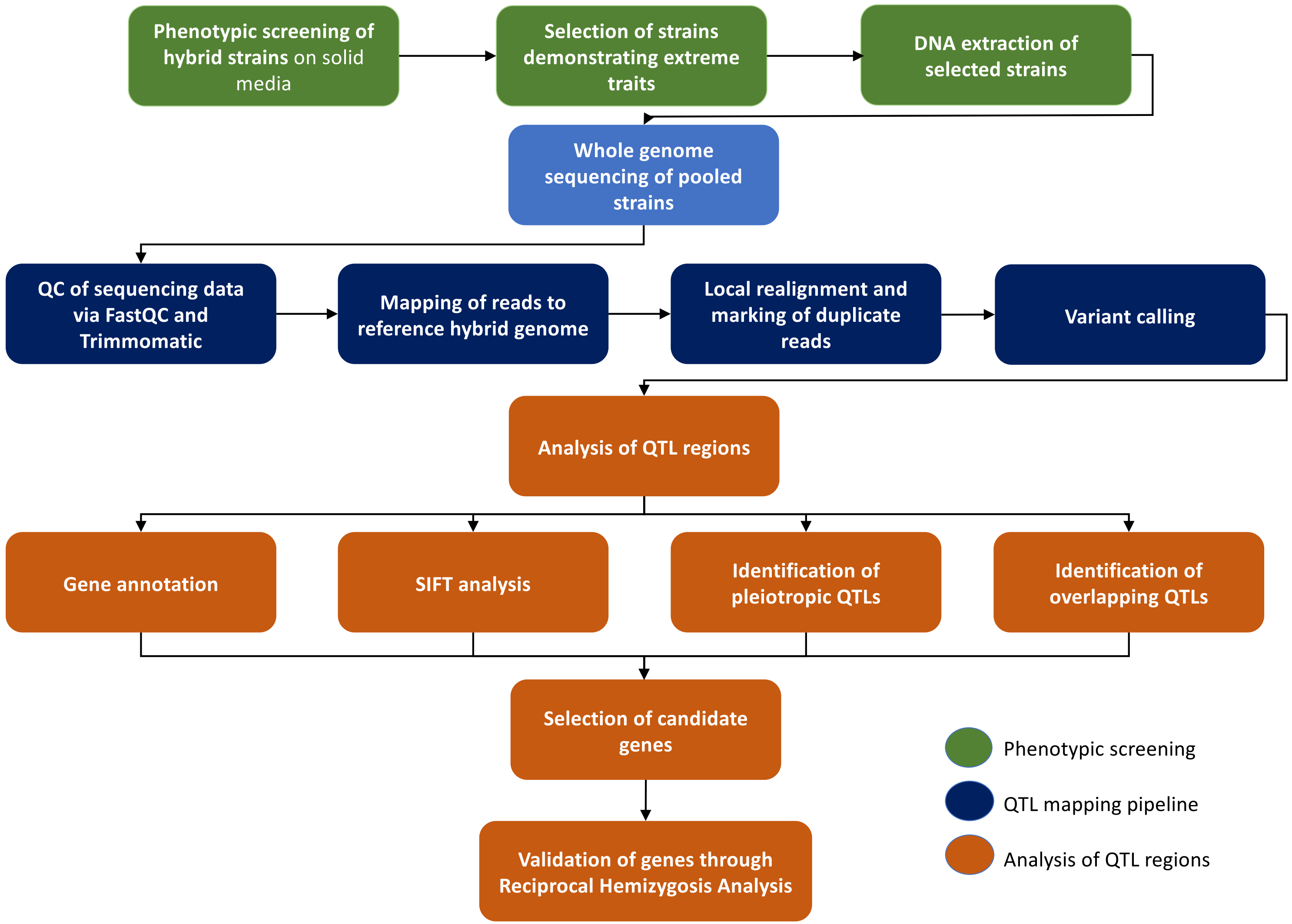


**Figure S1: Diagram representing each step of the in-house analysis pipeline to establish antifungal resistance QTLs in Saccharomyces hybrids.** A high-throughput phenotypic screening in solid media of a library of *S. cerevisiae x S. kudriavzevii* hybrids will allow the selection of a pool of the 20 best and the 20 worst hybrid strains in each condition tested. QTL analysis will allow to map the genetic regions underlying the phenotypic differences between the two pool and the data generated will be mined for the identification of candidate genes for further validation *in vivo*.

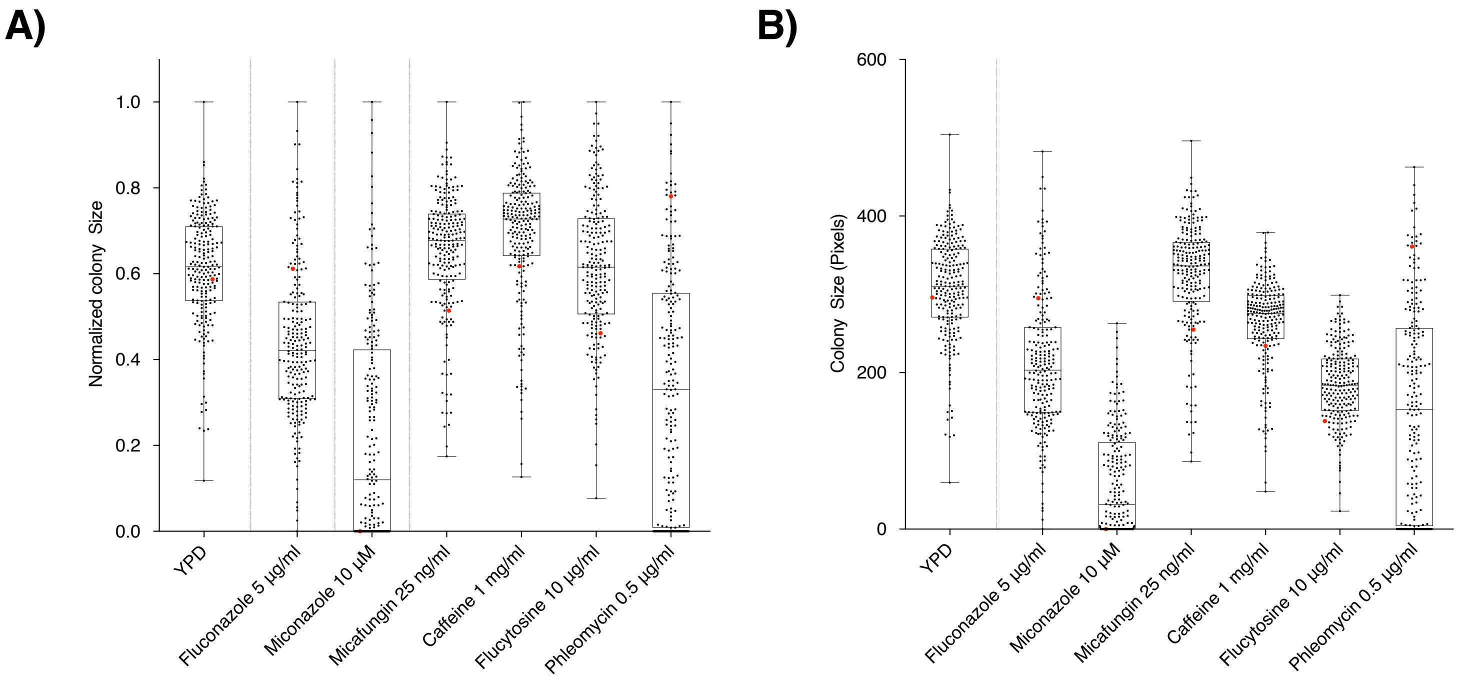


**Figure S2. Box plot of the fitness, in lower concentrations of drugs, of F12 diploid progeny for *S. cerevisiae/S. kudriavzevii*** hybrids after incubation with reduced concentrations of antifungal drugs expressed as colony size, (A) and as normalized colony size (B) parental tetraploid was used as a control and is highlighted in red. Each black dot represents a distinct F12 hybrid progeny. The upper and lower bars show the maximum and minimum values, the box represents the second and third quartile with the central line the median.

**Table S1 Descriptive statistics for phenotypic screening across YPD, YPD + Fluconazole (FCZ), YPD + miconazole (MCZ), YPD + caffeine (CAF), YPD micafungin (MCF), YPD + flucytosine (FCY) and YPD + phleomycin (BLE).** A total of 228 spores were plated along with parental control samples.

| | YPD | FCZ 5 µg/ml | FCZ 10 µg/ml | MCZ 10 µM | MCZ 20 µM | CAF 1 mg/ml | CAF 2.5 mg/ml | MCF 25 ng/ml | MCF 50 ng/ml | FCY 10 µg/ml | FCY 20 µg/ml |
|---|---|---|---|---|---|---|---|---|---|---|---|
| **25% Percentile** | 270.6 | 149 | 42.13 | 0 | 0 | 243.1 | 127.3 | 290.8 | 296 | 151.3 | 132.1 |
| **Median** | 310.5 | 203.3 | 87 | 31.5 | 0 | 275.8 | 168 | 336.3 | 351.3 | 184 | 158.3 |
| **wt colony size** | 296 | 295 | 172 | 0 | 0 | 234 | 101 | 255 | 224 | 138 | 131 |
| **75% Percentile** | 358 | 258 | 145.9 | 111.4 | 0 | 298.9 | 198.4 | 366.9 | 380.9 | 217.9 | 183.9 |
| **IQR** | 0.14 | 0.27 | 0.55 | 1.00 | 0.00 | 0.10 | 0.22 | 0.12 | 0.13 | 0.18 | 0.16 |
| **Range** | 444.5 | 482.5 | 254.5 | 263 | 294 | 331 | 269 | 409.5 | 468 | 276 | 222.5 |
| **Viability** | 100.0% | 99.6% | 87.7% | 64.5% | 23.2% | 100.0% | 99.6% | 100.0% | 100.0% | 100.0% | 100.0% |
| **Parental tetraploid control performance (position)** | 139 | 41 | 35 | 158 | Dead | 183 | 193 | 198 | 209 | 192 | 174 |

**Table S2 Genes identified in SIFT analysis as harbouring non-synonymous SNPs that are tolerated or deleterious to protein function.**

| Gene ID | Gene name | SIFT result |
|---|---|---|
| YDR132C | *MRX16* | TOLERATED |
| YDR135C | *YCF1* | DELETERIOUS |
| YDR137W | *RGP1* | TOLERATED |
| YDR138W | *HPR1* | TOLERATED |
| YDR140W | *MTQ2* | DELETERIOUS |
| YDR142C | *PEX7* | DELETERIOUS |
| YDR143C | *SAN1* | DELETERIOUS |
| YDR145W | *TAF12* | TOLERATED |
| YDR160W | *SSY1* | TOLERATED |
| YDR161W | *ACL4* | TOLERATED |
| **YDR162C** | *NBP2* | DELETERIOUS |
| YDR164C | *SEC1* | TOLERATED |
| YDR190C | *RVB1* | DELETERIOUS |
| YDR191W | *HST4* | TOLERATED |
| YDR192C | *NUP42* | TOLERATED |
| YDR195W | *REF2* | TOLERATED |
| YDR197W | *CBS2* | TOLERATED |
| YDR198C | *RKM2* | TOLERATED |
| YDR200C | *VPS64* | TOLERATED |
| YDR205W | *MSC2* | TOLERATED |
| YDR206W | *EBS1* | DELETERIOUS |
| **YDR207C** | *UME6* | DELETERIOUS |
| YDR208W | *MSS4* | DELETERIOUS |
| YDR211W | *GCD6* | TOLERATED |
| YDR212W | *TCP1* | TOLERATED |
| **YDR213W** | *UPC2* | DELETERIOUS |
| **YDR216W** | *ADR1* | DELETERIOUS |
| YDR217C | *RAD9* | DELETERIOUS |
| YDR218C | *SPR28* | DELETERIOUS |
| YDR219C | *MFB1* | TOLERATED |
| YER098W | *UBP9* | DELETERIOUS |
| YER100W | *UBC6* | TOLERATED |
| YER101C | *AST2* | TOLERATED |
| YER103W | *SSA4* | DELETERIOUS |
| YER130C | *COM2* | TOLERATED |
| YER132C | *PMD1* | TOLERATED |
| YER134C | | TOLERATED |
| YER165W | | DELETERIOUS |
| YER166W | | DELETERIOUS |
| YFL001W | *DEG1* | DELETERIOUS |

| | | |
|---|---|---|
| YFL002C | *SPB4* | TOLERATED |
| YFL003C | *MSH4* | TOLERATED |
| YFL004W | *VTC2* | DELETERIOUS |
| YFL007W | *BLM10* | TOLERATED |
| YFL008W | *SMC1* | TOLERATED |
| YFR001W | *LOC1* | DELETERIOUS |
| YFR003C | *YPI1* | TOLERATED |
| YFR005C | *SAD1* | TOLERATED |
| YFR006W | | TOLERATED |
| YFR007W | *YFH7* | TOLERATED |
| YFR010W | *UBP6* | TOLERATED |
| YGL044C | *RNA15* | TOLERATED |
| YGL045W | *RIM8* | TOLERATED |
| YGL048C | *RPT6* | TOLERATED |
| YKL008C | *LAC1* | TOLERATED |
| YML002W | | TOLERATED |
| YML004C | *GLO1* | TOLERATED |
| YML006C | *GIS4* | TOLERATED |
| YML007W | *YAP1* | TOLERATED |
| YML011C | *RAD33* | TOLERATED |
| YML013W | *UBX2* | DELETERIOUS |
| YML014W | *TRM9* | TOLERATED |
| YML016C | *PPZ1* | TOLERATED |
| YML017W | *PSP2* | TOLERATED |
| YML018C | | TOLERATED |
| YML020W | | DELETERIOUS |
| YML021C | *UNG1* | DELETERIOUS |
| YML068W | *ITT1* | TOLERATED |
| YML069W | *POB3* | TOLERATED |
| YMR004W | *MVP1* | TOLERATED |
| YMR005W | *TAF4* | TOLERATED |
| YMR006C | *PLB2* | TOLERATED |
| YMR008C | *PLB1* | TOLERATED |
| YMR011W | *HXT2* | TOLERATED |
| YMR013C | *SEC59* | DELETERIOUS |
| YMR185W | *RTP1* | DELETERIOUS |
| YMR186W | *HSC82* | TOLERATED |
| YMR187C | | TOLERATED |
| YMR189W | *GCV2* | TOLERATED |
| YMR190C | *SGS1* | DELETERIOUS |
| YMR191W | *SPG5* | TOLERATED |
| YMR194C-B | *CMC4* | TOLERATED |
| YMR196W | | TOLERATED |

| | | |
|---|---|---|
| YMR197C | *VTI1* | TOLERATED |
| **YNR069C** | *BSC5* | DELETERIOUS |
| YOR066W | *MSA1* | TOLERATED |
| YOR067C | *ALG8* | TOLERATED |
| YOR069W | *VPS5* | TOLERATED |
| YOR070C | *GYP1* | TOLERATED |
| YOR071C | *NRT1* | TOLERATED |
| YOR134W | *BAG7* | DELETERIOUS |
| YOR137C | *SIA1* | TOLERATED |
| YOR138C | *RUP1* | TOLERATED |
| **YOR140W** | *SFL1* | DELETERIOUS |
| YOR142W | *LSC1* | TOLERATED |
| YOR144C | *ELG1* | DELETERIOUS |
| YOR145C | *PNO1* | TOLERATED |
| YOR278W | *HEM4* | TOLERATED |
| YOR279C | *RFM1* | TOLERATED |
| YOR283W | | TOLERATED |
| YOR284W | *HUA2* | DELETERIOUS |
| YOR287C | *RRP36* | TOLERATED |
| YOR288C | *MPD1* | DELETERIOUS |
| YOR290C | *SNF2* | TOLERATED |
| YOR291W | *YPK9* | TOLERATED |
| YOR296W | | TOLERATED |
| YOR301W | *RAX1* | TOLERATED |
| YOR303W | *CPA1* | DELETERIOUS |
| YOR307C | *SLY41* | TOLERATED |
| YOR308C | *SNU66* | TOLERATED |
| YOR316C | *COT1* | DELETERIOUS |
| YOR317W | *FAA1* | DELETERIOUS |
| YOR319W | *HSH49* | DELETERIOUS |
| YOR320C | *GNT1* | TOLERATED |
| YOR321W | *PMT3* | TOLERATED |
| YOR323C | *PRO2* | DELETERIOUS |
| YOR355W | *GDS1* | DELETERIOUS |
| YOR356W | *CIR2* | TOLERATED |
| YOR359W | *VTS1* | TOLERATED |
| YOR363C | *PIP2* | DELETERIOUS |
| YOR371C | *GPB1* | TOLERATED |
| YOR372C | *NDD1* | TOLERATED |
| YBR072W | *HSP26* | TOLERATED |
| YBR074W | *PFF1* | TOLERATED |
| YBR076W | *ECM8* | TOLERATED |
| YBR077C | *SLM4* | TOLERATED |

| | | |
|---|---|---|
| YCR091W | *KIN82* | DELETERIOUS |
| YCR092C | *MSH3* | DELETERIOUS |
| YCR093W | *CDC39* | DELETERIOUS |
| YCR095C | *OCA4* | TOLERATED |
| YDR301W | *CFT1* | TOLERATED |
| YDR302W | *GPI11* | TOLERATED |
| YDR306C | *PFU1* | TOLERATED |
| YDR352W | *YPQ2* | DELETERIOUS |
| YDR356W | *SPC110* | DELETERIOUS |
| YDR359C | *EAF1* | DELETERIOUS |
| YDR361C | *BCP1* | TOLERATED |
| YDR362C | *TFC6* | TOLERATED |
| YDR363W | *ESC2* | TOLERATED |
| YDR364C | *CDC40* | DELETERIOUS |
| YDR365C | *ESF1* | TOLERATED |
| YDR367W | *KEI1* | TOLERATED |
| YDR368W | *YPR1* | TOLERATED |
| **YDR369C** | *XRS2* | TOLERATED |
| YDR372C | *VPS74* | TOLERATED |
| YDR374C | *PHO92* | TOLERATED |
| YDR376W | *ARH1* | DELETERIOUS |
| YDR379W | *RGA2* | DELETERIOUS |
| YDR380W | *ARO10* | TOLERATED |
| YDR383C | *NKP1* | TOLERATED |
| YDR402C | *DIT2* | TOLERATED |
| YDR405W | *MRP20* | TOLERATED |
| **YDR406W** | *PDR15* | DELETERIOUS |
| YDR407C | *TRS120* | TOLERATED |
| YDR408C | *ADE8* | TOLERATED |
| YDR409W | *SIZ1* | DELETERIOUS |
| YDR411C | *DFM1* | TOLERATED |
| YDR415C | | DELETERIOUS |
| YDR416W | *SYF1* | TOLERATED |
| YER083C | *GET2* | DELETERIOUS |
| YER085C | | TOLERATED |
| YER167W | *BCK2* | TOLERATED |
| YER169W | *RPH1* | TOLERATED |
| YGL063W | *PUS2* | TOLERATED |
| YGL141W | *HUL5* | TOLERATED |
| YGL142C | *GPI10* | TOLERATED |
| YGL143C | *MRF1* | TOLERATED |
| YGL144C | *ROG1* | DELETERIOUS |
| YGL145W | *TIP20* | TOLERATED |

| | | |
|---|---|---|
| YGL169W | *SUA5* | TOLERATED |
| YGL171W | *ROK1* | DELETERIOUS |
| YGL172W | *NUP49* | TOLERATED |
| YGR071C | *ENV11* | TOLERATED |
| YIR003W | *AIM21* | TOLERATED |
| YIR004W | *DJP1* | DELETERIOUS |
| YIR006C | *PAN1* | DELETERIOUS |
| YKR022C | *NTR2* | TOLERATED |
| YKR023W | *RQT4* | TOLERATED |
| YKR024C | *DBP7* | TOLERATED |
| YKR027W | *BCH2* | TOLERATED |
| YLR303W | *MET17* | DELETERIOUS |
| YLR385C | *SWC7* | TOLERATED |
| YMR018W | *PEX9* | TOLERATED |
| YMR020W | *FMS1* | DELETERIOUS |
| YMR021C | *MAC1* | TOLERATED |
| YMR023C | *MSS1* | DELETERIOUS |
| YMR114C | | TOLERATED |
| YMR115W | *MGR3* | DELETERIOUS |
| YMR163C | *INP2* | TOLERATED |
| YMR165C | *PAH1* | TOLERATED |
| **YMR167W** | *MLH1* | TOLERATED |
| YMR168C | *CEP3* | TOLERATED |
| YMR169C | *ALD3* | TOLERATED |
| YMR170C | *ALD2* | TOLERATED |
| **YNL102W** | *POL1* | TOLERATED |
| YNL103W | *MET4* | DELETERIOUS |
| YNL104C | *LEU4* | DELETERIOUS |
| YNR038W | *DBP6* | TOLERATED |
| YOR255W | *OSW1* | TOLERATED |
| YOR256C | *TRE2* | TOLERATED |
| YOR258W | *HNT3* | TOLERATED |
| YOR260W | *GCD1* | TOLERATED |
| YOR264W | *DSE3* | TOLERATED |
| YPL096W | *PNG1* | TOLERATED |
| YPL098C | *MGR2* | TOLERATED |
| YPL099C | *INA17* | TOLERATED |
| YPL100W | *ATG21* | DELETERIOUS |
| YPL101W | *ELP4* | TOLERATED |
| YPL126W | *NAN1* | TOLERATED |
| YPL130W | *SPO19* | DELETERIOUS |
| YPL131W | *RPL5* | DELETERIOUS |
| YPL133C | *RDS2* | DELETERIOUS |

| | | |
|---|---|---|
| YPR167C | *MET16* | TOLERATED |
| YDR259C | *YAP6* | TOLERATED |
| YDR261C | *EXG2* | TOLERATED |
| YDR262W | | TOLERATED |
| YDR263C | *DIN7* | TOLERATED |
| YDR265W | *PEX10* | TOLERATED |
| YDR266C | *HEL2* | TOLERATED |
| YDR311W | *TFB1* | TOLERATED |
| YDR312W | *SSF2* | TOLERATED |
| YDR313C | *PIB1* | TOLERATED |
| YDR314C | *RAD34* | TOLERATED |
| **YDR315C** | *IPK1* | TOLERATED |
| YDR316W | *OMS1* | TOLERATED |
| YDR317W | *HIM1* | TOLERATED |
| YDR318W | *MCM21* | DELETERIOUS |
| YDR319C | *YFT2* | DELETERIOUS |
| YDR320C | *SWA2* | DELETERIOUS |
| YDR321W | *ASP1* | TOLERATED |
| YJR036C | *HUL4* | TOLERATED |
| YJR062C | *NTA1* | TOLERATED |
| **YJR066W** | *TOR1* | TOLERATED |
| YJR067C | *YAE1* | TOLERATED |
| YJR107W | *LIH1* | TOLERATED |
| YJR109C | *CPA2* | TOLERATED |
| YKR003W | *OSH6* | TOLERATED |
| YKR004C | *ECM9* | TOLERATED |
| YKR007W | *MEH1* | TOLERATED |
| YKR009C | *FOX2* | TOLERATED |
| YKR011C | | TOLERATED |
| YKR015C | | DELETERIOUS |
| YKR016W | *MIC60* | TOLERATED |
| YLR381W | *CTF3* | DELETERIOUS |
| YLR382C | *NAM2* | DELETERIOUS |
| YOR108W | *LEU9* | TOLERATED |
| YOR109W | *INP53* | DELETERIOUS |
| YOR111W | | TOLERATED |
| YOR112W | *CEX1* | TOLERATED |
| YPL070W | *MUK1* | TOLERATED |
| YPL071C | | TOLERATED |
| YPL072W | *UBP16* | DELETERIOUS |
| YPL074W | *YTA6* | TOLERATED |
| YPL075W | *GCR1* | DELETERIOUS |
| YPL076W | *GPI2* | TOLERATED |

| | | |
|---|---|---|
| YPL082C | *MOT1* | TOLERATED |
| YPL083C | *SEN54* | TOLERATED |
| YPL084W | *BRO1* | TOLERATED |
| YPR011C | *MRX21* | TOLERATED |
| YPR013C | *CMR3* | DELETERIOUS |
| YPR016C | *TIF6* | TOLERATED |
| YPR017C | *DSS4* | DELETERIOUS |
| YPR018W | *RLF2* | TOLERATED |
| YPR022C | *SDD4* | TOLERATED |
| YPR025C | *CCL1* | TOLERATED |

**Table S3 List of potential causal genes identified in the S. cerevisiae genome of S. cerevisiae x S. kudriavzevii hybrids with non-synonymous SNPs predicted to be tolerated or deleterious by SIFT analysis**

| Condition | SIFT analysis | |
|---|---|---|
| | Tolerated | Deleterious |
| Fluconazole | *UBC6, COM2, LAC1, UBP6, MRX16, TAF12, HST4, NUP42, CBS2, RNA15, YAP1, PPZ1, PSP2, ITT1, HSC82, MSA1, ALG8, GYP1, HEM4, SLY41, SNU66, VTS1, NDD1, BCK2* | *DEG1, UPC2, ADR1, RAD9, DNF1, UBX2, SGS1, FAA1* |
| Flucytosine | *RDH54, ESC2, XRS2, NKP1, ADE8, MLH1, HNT3, RDS2, BCK2, ENV11* | *DNF1, BCH2, PDR15, POL1* |
| Micafungin | *BCK2, MEH1, EXG2, IPK1, TOR1, YTA6* | *DNF1* |

**Table S4 All genes identified on the *S. cerevisiae* genome within support intervals of QTL regions above the LOD threshold of 5, with no previously recorded genes linked to antifungal drug resistance.**

| Condition | Chromosome | LOD score | Gene ID | Common gene name | SIFT |
|---|---|---|---|---|---|
| Flucytosine | chrXVI | 27.36 | YPL101W | *ELP4* | Tolerated |
| Flucytosine | chrXVI | 27.36 | YPL100W | *ATG21* | Tolerated |
| Flucytosine | chrXVI | 27.36 | YPL099C | *INA17* | Deleterious |
| Flucytosine | chrXVI | 27.36 | YPL098C | *MGR2* | Tolerated |
| Flucytosine | chrXVI | 27.36 | YPL097W | *MSY1* | - |
| Flucytosine | chrXVI | 27.36 | YPL096C-A | *ERI1* | - |
| Flucytosine | chrXVI | 27.36 | YPL096W | *PNG1* | Tolerated |
| Flucytosine | ChrV | 21.58 | YER083C | *GET2* | Deleterious |
| Flucytosine | ChrV | 21.58 | YER084W | Uncharacterised | - |
| Flucytosine | ChrV | 21.58 | YER085C | Uncharacterised | Tolerated |
| Flucytosine | ChrV | 21.58 | YER086W | *ILV1* | - |
| Flucytosine | ChrV | 21.58 | YER087W | *AIM10* | - |
| Flucytosine | ChrV | 21.58 | YER087C-B | *SBH1* | - |
| Flucytosine | ChrX | 19.77 | YJR072C | *NPA3* | - |
| Flucytosine | chrXIV | 19.34 | YNL138W | *SRV2* | - |
| Flucytosine | chrXIV | 19.02 | YNR034W-A | *EGO4* | - |
| Flucytosine | chrXIV | 19.02 | YNR035C | *ARC35* | - |
| Flucytosine | chrXIV | 19.02 | YNR036C | *MRPS12* | - |
| Flucytosine | chrXIV | 19.02 | YNR037C | *RSM19* | - |
| Flucytosine | chrXVI | 16.43 | YPR165W | *RHO1* | Tolerated |
| Flucytosine | chrXVI | 16.43 | YPR166C | *MRP2* | - |
| Flucytosine | chrXVI | 16.43 | YPR167C | *MET16* | - |
| Flucytosine | chrXVI | 16.43 | YPR168W | *NUT2* | Tolerated |
| Flucytosine | chrXVI | 16.43 | YPR169W | *JIP5* | - |
| Flucytosine | ChrIX | 15.48 | YIR003W | *AIM21* | Deleterious |
| Flucytosine | ChrIX | 15.48 | YIR004W | *DJP1* | Deleterious |
| Flucytosine | ChrIX | 15.48 | YIR005W | *IST3* | - |
| Flucytosine | ChrIX | 15.48 | YIR006C | *PAN1* | Tolerated |
| Flucytosine | chrXV | 13.1 | YOR142W | *LSC1* | Tolerated |
| Flucytosine | chrXV | 13.1 | YOR143C | *THI80* | - |
| Flucytosine | chrXV | 13.1 | YOR144C | *ELG1* | Deleterious |
| Flucytosine | chrXV | 13.1 | YOR145C | *PNO1* | Tolerated |

| | | | | | |
|---|---|---|---|---|---|
| Flucytosine | ChrXII | 11.15 | YLR384C | *IKI3* | - |
| Flucytosine | ChrXII | 11.15 | YLR385C | *SWC7* | Tolerated |
| Flucytosine | ChrVII | 11.06 | YGL172W | *NUP49* | Tolerated |
| Flucytosine | ChrVII | 11.06 | YGL171W | *ROK1* | Deleterious |
| Flucytosine | ChrVII | 11.06 | YGL170C | *SPO74* | Deleterious |
| Flucytosine | ChrVII | 11.06 | YGL169W | *SUA5* | Tolerated |
| Flucytosine | ChrVII | 11.02 | YGL063W | *PUS2* | - |
| Flucytosine | chrXIII | 10.95 | YMR018W | *PEX9* | Deleterious |
| Flucytosine | chrXIII | 10.95 | YMR019W | *STB4* | Deleterious |
| Flucytosine | chrXIII | 10.95 | YMR020W | *FMS1* | Tolerated |
| Flucytosine | chrXIII | 10.95 | YMR021C | *MAC1* | Deleterious |
| Flucytosine | chrXIII | 10.95 | YMR022W | *UBC7* | - |
| Flucytosine | chrXIII | 10.95 | YMR023C | *MSS1* | Tolerated |
| Flucytosine | ChrIII | 8.31 | YCR090C | Uncharacterised | - |
| Flucytosine | ChrIII | 8.31 | YCR091W | *KIN82* | Tolerated |
| Flucytosine | ChrIII | 8.31 | YCR092C | *MSH3* | Tolerated |
| Flucytosine | ChrIII | 8.31 | YCR093W | *CDC39* | Deleterious |
| Flucytosine | ChrIII | 8.31 | YCR094W | *CDC50* | - |
| Flucytosine | ChrIII | 8.31 | YCR095C | *OCA4* | Tolerated |
| Flucytosine | ChrIII | 8.31 | YCR096C | *HMRA2* | - |
| Flucytosine | ChrIII | 8.31 | YCR097W | *HMRA1* | - |
| Micafungin | chrX | 10.15 | YJR036C | *HUL4* | Tolerated |
| Micafungin | chrXV | 9.06 | YOR108W | *LEU9* | Tolerated |
| Micafungin | chrXV | 9.06 | YOR109W | *INP53* | Deleterious |
| Micafungin | chrXV | 9.06 | YOR110W | *TFC7* | - |
| Micafungin | chrXV | 9.06 | YOR111W | Uncharacterised | Tolerated |
| Micafungin | chrXV | 9.06 | YOR112W | *CEX1* | Tolerated |
| Micafungin | chrIV | 8.62 | YDR144C | *MKC7* | - |
| Micafungin | chrIV | 8.62 | YDR145W | *TAF12* | Tolerated |
| Micafungin | chrIV | 8.62 | YDR146C | *SWI5* | - |
| Micafungin | chrIV | 8.62 | YDR147W | *EKI1* | - |
| Micafungin | chrIV | 8.62 | YDR148C | *KGD2* | - |
| Micafungin | chrX | 7.72 | YJR107W | *LIH1* | Tolerated |
| Micafungin | chrX | 7.72 | YJR108W | *ABM1* | - |
| Micafungin | chrX | 7.72 | YJR109C | *CPA2* | Tolerated |
| Micafungin | chrXII | 6.96 | YLR380W | *CSR1* | - |

| | | | | | |
|---|---|---|---|---|---|
| Micafungin | chrXII | 6.96 | YLR381W | *CTF3* | Deleterious |
| Micafungin | chrXII | 6.96 | YLR382C | *NAM2* | Deleterious |
| Micafungin | chrXVI | 6.76 | YPR009W | *SUT2* | - |
| Micafungin | chrXVI | 6.76 | YPR010C | *RPA135* | - |
| Micafungin | chrXVI | 6.76 | YPR010C-A | *MIN8* | - |
| Micafungin | chrXVI | 6.76 | YPR011C | *MRX21* | Tolerated |
| Micafungin | chrXVI | 6.76 | YPR013C | *CMR3* | Deleterious |
| Micafungin | chrXVI | 6.76 | YPR015C | Uncharacterised | - |
| Micafungin | chrXVI | 6.76 | YPR016C | *TIF6* | Tolerated |
| Micafungin | chrXVI | 6.76 | YPR017C | *DSS4* | Deleterious |
| Micafungin | chrXVI | 6.76 | YPR018W | *RLF2* | Tolerated |
| Micafungin | chrXVI | 6.76 | YPR019W | *MCM4* | - |
| Micafungin | chrXVI | 6.76 | YPR020W | *ATP20* | - |
| Micafungin | chrXVI | 6.76 | YPR021C | *AGC1* | - |
| Micafungin | chrXVI | 6.76 | YPR022C | *SDD4* | Tolerated |
| Micafungin | chrXVI | 6.76 | YPR023C | *EAF3* | - |
| Micafungin | chrXVI | 6.76 | YPR024W | *YME1* | - |
| Micafungin | chrXVI | 6.76 | YPR025C | *CCL1* | Tolerated |
| Fluconazole | chrIV | 16.73 | YDR164C | *SEC1* | Tolerated |
| Fluconazole | chrIV | 16.73 | YDR163W | *CWC15* | - |
| Fluconazole | chrIV | 16.73 | YDR162C | *NBP2* | Deleterious |
| Fluconazole | chrIV | 16.73 | YDR161W | *ACL4* | Tolerated |
| Fluconazole | chrIV | 16.73 | YDR160W | *SSY1* | Tolerated |

**Table S5 All genes identified on the *S. kudrivzevii* genome within support intervals of QTL regions above the LOD threshold of 5, with no previously recorded genes linked to antifungal drug resistance.**

| Condition | Chromosome | LOD score | Gene ID | Common gene name |
|---|---|---|---|---|
| Flucytosine | Skud_4 | 29.58 | YDR169C | *STB3* |
| Flucytosine | Skud_4 | 29.58 | YDR170C | *SEC7* |
| Flucytosine | Skud_4 | 29.58 | SKUD0Dtrna7Q | |
| Flucytosine | Skud_11 | 28.47 | YKL181W | *PRS1* |
| Flucytosine | Skud_11 | 28.47 | SKUD0K00470 | |
| Flucytosine | Skud_11 | 28.47 | YKL179C | *COY1* |
| Flucytosine | Skud_12 | 21.19 | SKUD0Ltrna6R | |
| Flucytosine | Skud_12 | 21.19 | YLR114C | *AVL9* |
| Flucytosine | Skud_12 | 21.19 | YLR115W | *CFT2* |
| Flucytosine | Skud_12 | 21.19 | YLR116W | *MSL5* |
| Flucytosine | Skud_12 | 21.19 | YLR117C | *CLF1* |
| Flucytosine | Skud_12 | 21.19 | YLR118C | *TML25* |
| Flucytosine | Skud_12 | 21.19 | YLR119W | *SRN2* |
| Flucytosine | Skud_12 | 21.19 | YLR120C | *YPS1* |
| Flucytosine | Skud_12 | 21.19 | YLR121C | *YPS3* |
| Flucytosine | Skud_12 | 21.19 | SKUD0L01780 | |
| Flucytosine | Skud_12 | 21.19 | YLR125W | |
| Flucytosine | Skud_12 | 21.19 | YLR126C | |
| Flucytosine | Skud_12 | 21.19 | YLR127C | *APC2* |
| Flucytosine | Skud_12 | 21.19 | YLR128W | *DCN1* |
| Flucytosine | Skud_12 | 21.19 | YLR129W | *DIP2* |
| Flucytosine | Skud_12 | 21.19 | YLR130C | *ZRT2* |
| Flucytosine | Skud_12 | 21.19 | YLR131C | *ACE2* |
| Flucytosine | Skud_12 | 21.19 | YLR132C | *USB1* |
| Flucytosine | Skud_12 | 21.19 | YLR133W | *CKI1* |
| Flucytosine | Skud_12 | 21.19 | YLR134W | *PDC5* |
| Flucytosine | Skud_12 | 21.19 | YLR135W | *SLX4* |
| Flucytosine | Skud_12 | 21.19 | YLR136C | *TIS11* |
| Flucytosine | Skud_12 | 21.19 | YLR137W | *RKM5* |
| Flucytosine | Skud_12 | 21.19 | YLR138W | *NHA1* |
| Flucytosine | Skud_12 | 21.19 | YLR139C | *SLS1* |
| Flucytosine | Skud_2 | 20.26 | YBR150C | *TBS1* |
| Flucytosine | Skud_2 | 20.26 | YBR151W | *APD1* |
| Flucytosine | Skud_4 | 18.5 | YDR306C | *PFU1* |
| Flucytosine | Skud_4 | 18.5 | SKUD0Dtrna10V | |
| Flucytosine | Skud_4 | 18.5 | YDR307W | *PMT7* |
| Flucytosine | Skud_4 | 18.5 | YDR308C | *SRB7* |

| | | | | |
|---|---|---|---|---|
| Flucytosine | Skud_4 | 18.5 | YDR309C | *GIC2* |
| Flucytosine | Skud_8 | 18.08 | YHR176W | *FMO1* |
| Flucytosine | Skud_8 | 18.08 | YHR177W | *ROF1* |
| Flucytosine | Skud_7 | 17.65 | YGR243W | *MPC3* |
| Flucytosine | Skud_7 | 17.65 | YGR244C | *LSC2* |
| Flucytosine | Skud_7 | 17.65 | YGR245C | *SDA1* |
| Flucytosine | Skud_7 | 17.65 | YGR246C | *BRF1* |
| Flucytosine | Skud_7 | 17.65 | YGR247W | *CPD1* |
| Flucytosine | Skud_7 | 17.65 | YGR248W | *SOL4* |
| Flucytosine | Skud_11 | 14.17 | YKL104C | *GFA1* |
| Flucytosine | Skud_11 | 14.17 | YKL103C | *APE1* |
| Flucytosine | Skud_16 | 13.13 | YPL010W | *RET3* |
| Flucytosine | Skud_16 | 13.13 | YPL009C | *RQC2* |
| Flucytosine | Skud_16 | 13.13 | YPL008W | *CHL1* |
| Flucytosine | Skud_16 | 13.13 | YPL007C | *TFC8* |
| Flucytosine | Skud_16 | 13.13 | YPL006W | *NCR1* |
| Flucytosine | Skud_16 | 13.13 | YPL005W | *AEP3* |
| Flucytosine | Skud_16 | 13.13 | YPL004C | *LSP1* |
| Flucytosine | Skud_16 | 13.13 | YPL003W | *ULA1* |
| Flucytosine | Skud_16 | 13.13 | YPL002C | *SNF8* |
| Flucytosine | Skud_16 | 13.13 | YPL001W | *HAT1* |
| Flucytosine | Skud_14 | 12.08 | YNL153C | *GIM3* |
| Flucytosine | Skud_14 | 12.08 | YNL152W | *INN1* |
| Flucytosine | Skud_14 | 12.08 | YNL151C | *RPC31* |
| Flucytosine | Skud_14 | 12.08 | YNL149C | *PGA2* |
| Flucytosine | Skud_14 | 12.08 | YNL148C | *ALF1* |
| Flucytosine | Skud_14 | 12.08 | YNL147W | *LSM7* |
| Flucytosine | Skud_14 | 12.08 | YNL146W | |
| Flucytosine | Skud_14 | 12.08 | SKUD0N01810 | |
| Flucytosine | Skud_14 | 12.08 | YNL144C | |
| Flucytosine | Skud_11 | 8.44 | YKL068W-A | |
| Flucytosine | Skud_11 | 8.44 | YKL068W | *NUP100* |
| Flucytosine | Skud_11 | 8.44 | SKUD0Ktrna1H | |
| Flucytosine | Skud_11 | 8.44 | YKL067W | *YNK1* |
| Flucytosine | Skud_11 | 8.44 | YKL065W-A | *DPC7* |
| Flucytosine | Skud_11 | 8.44 | YKL065C | *YET1* |
| Flucytosine | Skud_11 | 8.44 | YKL064W | *MNR2* |
| Flucytosine | Skud_11 | 8.44 | YKL063C | |
| Flucytosine | Skud_11 | 8.44 | SKUD0K01520 | |
| Flucytosine | Skud_11 | 8.44 | YKL062W | *MSN4* |
| Flucytosine | Skud_11 | 8.44 | YKL061W | *BLI1* |
| Flucytosine | Skud_11 | 8.44 | YKL060C | *FBA1* |
| Flucytosine | Skud_11 | 8.44 | YKL059C | *MPE1* |

| | | | | |
|---|---|---|---|---|
| Flucytosine | Skud_11 | 8.44 | YKL058W | *TOA2* |
| Flucytosine | Skud_11 | 8.44 | YKL057C | *NUP120* |
| Flucytosine | Skud_11 | 8.44 | YKL056C | *TMA19* |
| Flucytosine | Skud_11 | 8.39 | YKR079C | *TRZ1* |
| Flucytosine | Skud_11 | 8.39 | YKR080W | *MTD1* |
| Flucytosine | Skud_11 | 8.39 | YKR081C | *RPF2* |
| Flucytosine | Skud_11 | 8.39 | YKR082W | *NUP133* |
| Flucytosine | Skud_11 | 8.39 | YKR083C | *DAD2* |
| Flucytosine | Skud_11 | 8.39 | YKR084C | *HBS1* |
| Flucytosine | Skud_11 | 8.39 | YKR085C | *MRPL20* |
| Flucytosine | Skud_2 | 8.22 | YBR024W | *SCO2* |
| Flucytosine | Skud_2 | 8.22 | YBR025C | *OLA1* |
| Flucytosine | Skud_2 | 8.22 | YBR026C | *ETR1* |
| Flucytosine | Skud_2 | 8.22 | YBR028C | *YPK3* |
| Flucytosine | Skud_2 | 7.73 | YBR221C | *PDB1* |
| Flucytosine | Skud_2 | 7.73 | YBR222C | *PCS60* |
| Flucytosine | Skud_2 | 7.73 | YBR223C | *TDP1* |
| Flucytosine | Skud_2 | 7.73 | YBR225W | |
| Flucytosine | Skud_2 | 7.73 | YBR227C | *MCX1* |
| Flucytosine | Skud_2 | 7.73 | YBR228W | *SLX1* |
| Flucytosine | Skud_5 | 7.27 | YER156C | *MYG1* |
| Flucytosine | Skud_5 | 7.27 | YER157W | *COG3* |
| Flucytosine | Skud_5 | 7.27 | SKUD0Etrna9E | |
| Flucytosine | Skud_5 | 7.27 | YER158C | |
| Flucytosine | Skud_5 | 7.27 | YER159C | *BUR6* |
| Flucytosine | Skud_5 | 7.27 | SKUD0Etrna14R | |
| Flucytosine | Skud_5 | 7.27 | YER161C | *SPT2* |
| Flucytosine | Skud_5 | 7.27 | YER162C | *RAD4* |
| Flucytosine | Skud_5 | 7.27 | YER163C | *GCG1* |
| Flucytosine | Skud_10 | 6.82 | YJL019W | *MPS3* |
| Flucytosine | Skud_10 | 6.82 | YJL016W | *TPH3* |
| Flucytosine | Skud_10 | 6.82 | YJL014W | *CCT3* |
| Flucytosine | Skud_10 | 6.82 | YJL013C | *MAD3* |
| Flucytosine | Skud_10 | 6.82 | YJL012C | *VTC4* |
| Flucytosine | Skud_10 | 6.82 | YJL011C | *RPC17* |
| Flucytosine | Skud_10 | 6.82 | SKUD0Jtrna1K | |
| Flucytosine | Skud_10 | 6.82 | SKUD0Jtrna1W | |
| Micafungin | Skud_2 | 9.76 | YBR255C-A | *RCF3* |
| Micafungin | Skud_2 | 9.76 | YBR256C | *RIB5* |
| Micafungin | Skud_2 | 9.76 | YBR257W | *POP4* |
| Micafungin | Skud_2 | 9.76 | YBR258C | *SHG1* |
| Micafungin | Skud_2 | 9.76 | YBR259W | |
| Micafungin | Skud_11 | 11.28 | YKL171W | *NNK1* |

| | | | | |
|---|---|---|---|---|
| Micafungin | Skud_11 | 11.28 | YKL170W | *MRPL38* |
| Micafungin | Skud_11 | 11.28 | YKL168C | *KKQ8* |
| Micafungin | Skud_11 | 11.28 | YKL167C | *MRP49* |
| Micafungin | Skud_11 | 11.28 | YKL166C | *TPK3* |
| Micafungin | Skud_11 | 11.28 | YKL165C | *MCD4* |
| Micafungin | Skud_11 | 11.28 | SKUD0Ktrna1E | |
| Micafungin | Skud_11 | 11.28 | YKL164C | *PIR1* |
| Micafungin | Skud_11 | 17.13 | YKL100C | *YPF1* |
| Micafungin | Skud_11 | 17.13 | YKL099C | *UTP11* |
| Micafungin | Skud_11 | 17.13 | YKL098W | *MTC2* |
| Micafungin | Skud_11 | 17.13 | SKUD0K01210 | |
| Fluconazole | Skud_12 | 18.08 | YLR321C | *SFH1* |
| Fluconazole | Skud_12 | 18.08 | YLR323C | *CWC24* |
| Fluconazole | Skud_12 | 18.08 | YLR324W | *PEX30* |
| Fluconazole | Skud_12 | 18.08 | YLR325C | *RPL38* |
| Fluconazole | Skud_12 | 18.08 | YLR326W | |
| Fluconazole | Skud_12 | 18.08 | YLR327C | *TMA10* |
| Fluconazole | Skud_12 | 18.08 | SKUD0Ltrna4S | |
| Fluconazole | Skud_12 | 18.08 | YLR328W | *NMA1* |
| Fluconazole | Skud_12 | 18.08 | SKUD0L03620 | |
| Fluconazole | Skud_13 | 11.5 | YMR018W | *PEX9* |
| Fluconazole | Skud_13 | 11.5 | YMR019W | *STB4* |

**Table S6 List of strains generated for reciprocal hemizygosis analysis**

| **Hybrid** | **Strain Number** | **Deletion** |
|---|---|---|
| *S. cerevisiae x S. cerevisiae* | OS104 x OS253 | $BCK2^{OS104}$::kanMX<br>$BCK2^{OS253}$::natMX<br>$DNF1^{OS104}$::kanMX<br>$DNF1^{OS253}$::natMX |
| *S. cerevisiae x S. kudriavzevii* | (OS104 x IFO1802) x (OS253 x OS575) | $BCK2^{OS104}$::BLE<br>$BCK2^{OS253}$::BLE<br>$DNF1^{OS104}$::BLE<br>$DNF1^{OS253}$::BLE |

**Table S7 Values for growth dynamics of reciprocal hemizygotes for interspecies and intraspecies hybrids for the genes *DNF1* and *BCK2* in flucytosine and micafungin, respectively.** Values were calcluated using the growth curver R package and graphpad prism for statistical analysis. Shown are the means and standard error for each condition, and significant difference was calculated using the t-test (P<0.05).

| **Strain Background** | ***Sc/Sk$_m$ DNF1Δ*** | | |
|---|---|---|---|
| | **$Sc^{OS104/Δ}$** | **$Sc^{Δ/OS253}$** | **P value** |
| Specific growth rate $h^{-1}$ | 0.29±0.005 | 0.27±0.004 | 0.0011** |
| Maximum biomass ($OD_{600}$) | 1.40±0.005 | 1.33±0.003 | 0.0001*** |
| $T_{mid}$ | 11.31±0.057 | 11.96±0.070 | 0.0001*** |
| Integral area | 40.79±0.210 | 37.75±0.117 | 0.0001*** |
| **Strain Background** | **Sc/Sc *DNF1Δ*** | | |
| | **$Sc^{OS104/Δ}$** | **$Sc^{Δ/OS253}$** | **P value** |
| Specific growth rate $h^{-1}$ | 0.27±0.005 | 0.25±0.003 | 0.0692 |
| Maximum biomass ($OD_{600}$) | 1.22±0.006 | 1.225±0.007 | 0.7218 |
| $T_{mid}$ | 12.23±0.114 | 12.17±0.071 | 0.6544 |
| Integral area | 40.86±0.257 | 41.05±0.269 | 0.6271 |
| **Strain Background** | **Sc/Sk$_m$ *BCK2Δ*** | | |
| | **$Sc^{OS104/Δ}$** | **$Sc^{Δ/OS253}$** | **P value** |
| Specific growth rate $h^{-1}$ | 0.40±0.012 | 0.36±0.026 | 0.1819 |
| Maximum biomass ($OD_{600}$) | 1.34±0.027 | 1.38±0.043 | 0.4061 |
| $T_{mid}$ | 14.97± 0.613 | 18.83±0.587 | 0.0003*** |
| Integral area | 41.30±1.509 | 37.08±0.789 | 0.0289* |
| Strain Background | **Sc/Sc *BCK2Δ*** | | |
| | **$Sc^{OS104/Δ}$** | **$Sc^{Δ/OS253}$** | **P value** |
| Specific growth rate $h^{-1}$ | 0.42±0.015 | 0.37±0.020 | 0.0383* |
| Maximum biomass ($OD_{600}$) | 1.32±0.025 | 1.37±0.032 | 0.2509 |
| $T_{mid}$ | 16.15±0.770 | 17.19±0.961 | 0.4064 |
| Integral area | 39.38±1.580 | 39.09±1.204 | 0.8825 |

**Table S8: List of primers used in this study for the generation of reciprocal hemizygotes.**

| Primer name | Primer sequence |
|---|---|
| DNF1 Kan up | ATCCTTTATTTCCTCCTGGAAGTGAAATAAAAACGCAGTTAACTTTAAAAAGGAAAGTTAGTTGATAATTCGTACGCTGCAGGTCGAC |
| DNF1 Kan down | AATATTATGTAATATTATATATACAGTGAAAAGCGGAGTTCCATCAGCTATGGTTCGGATATCATTATTAATCGATGAATTCGAGCTCG |
| DNF1 conf A | CGCATTCAACTAACTGGAGG |
| DNF1 conf D | GCCAGTTAGTGGAGAGTTC |
| DNF1 conf B | GGGTTGCCTTCTTCATCTATC |
| DNF1 conf C | CAGCGCGTATCTATGGAG |
| DNF1 Ble Up | ATCCTTTATTTCCTCCTGGAAGTGAAATAAAAACGCAGTTAACTTTAAAAAGGAAAGTTAGTTGATAATTCTGTTTAGCTTGCCTCGTCC |
| DNF1 Ble Down | AATATTATGTAATATTATATATACAGTGAAAAGCGGAGTTCCATCAGCTATGGTTCGGATATCATTATTATGATGCGGTATTTTCTCCTTACG |
| BCK2 Kan up | AGAAAGCTCAGAAGGAGATCTAAGACAAGAAGCAAGGCAGTCTTGATTTATCGCTCAAAACCAATAGCACCCGTACGCTGCAGGTCGAC |
| BCK2 Kan down | CAGTTTTTAACTTTTTCGAAAGATATCTGTTACTATTTTTGAACTTTTTTTTTTTTTTTTCATTCCTTTTATCGATGAATTCGAGCTCG |
| BCK2 conf A | GCTGGGAACTCTCCACTAAC |
| BCK2 conf D | CTACTTCAAATGAAGCCAGGTC |
| BCK2 conf B | CTGGAGTGCTACATGAGC |
| BCK2 conf C | CGATGGTCATACTTTCGTTCC |
| BCK2 Ble Up | AGAAAGCTCAGAAGGAGATCTAAGACAAGAAGCAAGGCAGTCTTGATTTATCGCTCAAAACCAATAGCACCTGTTTAGCTTGCCTCGTCC |
| BCK2 Ble Down | CAGTTTTTAACTTTTTCGAAAGATATCTGTTACTATTTTTGAACTTTTTTTTTTTTTTTTCATTCCTTTTTGATGCGGTATTTTCTCCTTACG |
| Kan_ORF REV | AACGTGAGTCTTTTCCTTACC |
| Kan_ORF FWD | TCTCCTTCATTACAGAAACG |
| Nat_ORF REV | TTCGTCGTCCGATTCGTCGT |
| Nat_ORF FWD | TGACCGTCGAGGACATCGA |
| Ble_ORF REV | CTCATGAGATGCCTGCAAGC |
| Ble_ORF FWD | CCGCCGCCTTCTATGAAAG |
